# Protein-interaction network analysis reveals a role of Prp19 splicing factor in transcription of both intron-containing and intron-lacking genes

**DOI:** 10.1101/2025.03.31.646471

**Authors:** Katherine Dwyer, Mary-Ann Essak, Ahlam Awada, Zuzer Dhoondia, Athar Ansari

## Abstract

We have previously demonstrated that the transcription-dependent interaction of the promoter and terminator ends of a gene, which results in the formation of a gene loop, is facilitated by the interaction of the general transcription factor TFIIB with the CF1, CPF and Rat1 termination complexes. To further elucidate the protein-protein interactions that stabilize gene loop, we performed mass spectrometry of affinity purified termination complexes from chromatin fraction. Quantitative proteomic analysis revealed additional interactions of termination factors with TFIID and SAGA complex. Since gene looping of intron-containing genes involves additional contacts of the promoter and terminator with the intron, we examined if termination factors interact with the splicing factors as well. All three termination complexes displayed statistically significant interactions with Prp19, Prp43, Sub2, Snu114, Brr2 and Smb1 splicing factors. Since Prp43 and Prp19 consistently emerged as the interactor of both initiation and termination factors, we affinity-purified both and performed mass spectrometry. Prp19 exhibited interactions with subunits of TFIID, CPF complex, and the RSC chromatin remodeling complex. These interactions were observed exclusively in the chromatin context, thereby implicating the factor in transcription of protein coding genes. Since fewer than 4% of yeast genes contain introns, we hypothesized that Prp19 might have a broader role in RNAPII transcription cycle. Auxin-mediated depletion of Prp19 resulted in about two-fold decrease in transcription of a subset of both intron-containing and intron-lacking genes. Specifically, the promoter recruitment of TBP registered a significant decline in the absence of Prp19. Chromatin immunoprecipitation (ChIP) analysis revealed crosslinking of Prp19 to the promoter proximal as well as downstream regions of both intronic and non-intronic genes. These findings demonstrate that Prp19 has a novel role in the initiation step of transcription in yeast.

## Introduction

In the complex landscape of eukaryotic gene expression, the process of transcription by RNA polymerase II is intricately linked with the modification of nascent mRNA by 5’ capping, splicing, and cleavage-polyadenylation at the 3’ end (Proudfoot et al., 2002; Bentley, 2014). These processing steps of mRNA maturation predominantly occur cotranscriptionally. The different steps of transcription and cotranscriptional RNA processing occur in a coordinated fashion and are fine-tuned by a complex network of molecular interactions. One intriguing phenomenon that has garnered recent attention in eukaryotic transcription is gene looping. Gene looping represents a spatial arrangement where the chromosomal promoter and terminator ends of a gene engage in physical contact to form a looped structure (O’Sullivan et al., 2004; Ansari and Hampsey, 2005; Al-Husini et al., 2020). Evidence suggest that this architectural conformation may be crucial for coordinating various transcriptional and cotranscriptional events, including termination, termination-coupled reinitiation, and promoter directionality, thus playing a pivotal role in regulating both transcription and cotranscriptional mRNA processing (Tan-Wong et al., 2012; Al Husini et al., 2013; Grzechnik et al., 2014; Al-Husini et al., 2020). Gene loops are formed in a transcription-dependent manner and were first demonstrated in the yeast *Saccharomyces cerevisiae* (O’Sullivan et al., 2004; Ansari and Hampsey, 2005). Subsequent studies have documented the presence of gene looping in other eukaryotes (Henriques et al., 2012; Crevillen et al., 2013; Le May et al., 2012; Shibayama et al., 2014; Gangliardi et al., 2019; Zhao et al., 2021; Terrone et al., 2022).

Research in yeast has identified general transcription factors and termination complexes as crucial players in gene looping (O’Sullivan et al., 2004; Ansari and Hampsey, 2005; El Kaderi et al., 2009; Medler et al., 2011; Mukundan and Ansari, 2013; Al Husini et al., 2013; Medler and Ansari, 2015; Allepuz-Fuster et al., 2019; Terrone et al., 2022). Notably, the general transcription factor TFIIB has been implicated in stabilizing gene loops through its interaction with key termination complex components such as CF1, CPF, and Rat1 in yeast (Medler et al., 2011; O’Brien and Ansari, 2024). Intriguingly, a complex comprising the general transcription factor TFIID and 3’ end processing-termination factors has been purified from mammalian cells as well (Dantonel et al., 1997). In yeast, Mediator subunit Srb5 and TFIIH subunit Kin28 have also been implicated in gene loop stabilization (Mukundan and Ansari, 2013; Medler and Ansari, 2015). Recent studies in our laboratory conducted mass spectrometry analyses of affinity-purified initiation and termination factor complexes from chromatin fractions, to explore the protein-protein interactions underlying gene looping. These investigations revealed novel associations of TFIIB and Rat1 with splicing factors (Dhoondia et al., 2021; O’Brien and Ansari, 2024).

Gene looping of intron-containing genes requires additional contacts between the promoter, terminator, and the intronic regions during transcription (Moabbi et al., 2012). The observed interactions between TFIIB and splicing factors as well as between Rat1 and splicing factors, suggest a broader, yet currently unidentified, role of these molecular players in gene loop formation and consequently in transcription and cotranscriptional mRNA processing. Given the cotranscriptional nature of splicing, it is reasonable to hypothesize an interaction of splicing factors with components of the transcription machinery. This idea is further supported by known functional interactions between splicing factors and transcription machinery in higher eukaryotes (Robert et al., 2002; Kornblihtt et al., 2004; Rambout et al., 2017; Carnesecchi et al., 2022; Göös et al., 2022; Nabeel-Shah et al., 2024). In mammalian systems, over 90% of protein coding genes contain introns. The possibility of spliceosome-transcription machinery relationships is therefore not surprising. In yeast, there is genetic evidence of interactions between transcription initiation and termination factors and splicing factors (Costanzo et al., 2016; Shao et al., 2020). However, since less than 4% of yeast genes contain introns (Ares et al., 1999), it is tempting to speculate that splicing factor-transcription factor interactions play a broader, splicing-independent role in transcriptional regulation.

To explore this possibility, we set out to probe for novel interactions of termination factors with the splicing machinery. Our results revealed multiple interactions of all three yeast 3’ end processing complexes, CPF, CF1 and Rat1, with splicing factors. A reciprocal approach involving purification of the splicing factor Prp19 confirmed its interaction with both initiation and termination factors. Furthermore, Prp19 exhibited a strong interaction with the RSC chromatin remodeling complex. The observation that Prp19 shows statistically significant interactions with TFIID, CPF complex, and RSC complex exclusively within the chromatin context emphasizes the transcription-dependent nature of these interactions. These findings suggest a potential function of splicing factors in transcription of RNAPII-transcribed genes. Auxin-mediated depletion of Prp19 revealed its moderate stimulatory influence on the transcription of a subset of both intronic and non-intronic genes. Specifically, Prp19 crosslinked to the promoter-proximal region of genes and facilitated the recruitment of TFIID subunit TBP. These novel findings challenge the traditional view of splicing factors solely as mediators of RNA processing, positioning them as integral components in the regulation of transcription by RNA polymerase II.

## Results

Our previous work revealed that the interaction of TFIIB with termination factors is crucial for the promoter-terminator communication through gene looping and plays a pivotal role in transcription termination (O’Brien et al., 2024). Interestingly, both TFIIB and Rat1 also exhibit interactions with a number of splicing factors in the chromatin context (O’Brien and Ansari, 2024; Dhoondia et al., 2021). To further explore the possibility of novel protein-protein interactions that take place on chromatin during transcription, we purified all three termination complexes of yeast: CPF, CF1, and Rat1, from soluble and chromatin fractions of yeast cell lysate (Fig. 1A). Specifically, CPF was purified from a strain with Myc-tagged Ssu72, CF1 from a strain with HA-tagged Rna15, and Rat1 from a strain with HA-tagged Rat1 (Fig. 1B). Yeast cells were lysed, and the lysate was separated into soluble and chromatin fractions by differential centrifugation. Affinity chromatography was then employed to purify CPF, CF1, and Rat1 complexes from both fractions (Fig. 1A). Mass spectrometry was performed to identify interacting protein partners of the three complexes. The results revealed some expected interactions of termination factors with RNAPII and the general transcription factors, and some rather novel interactions with splicing factors.

**Figure 1:**
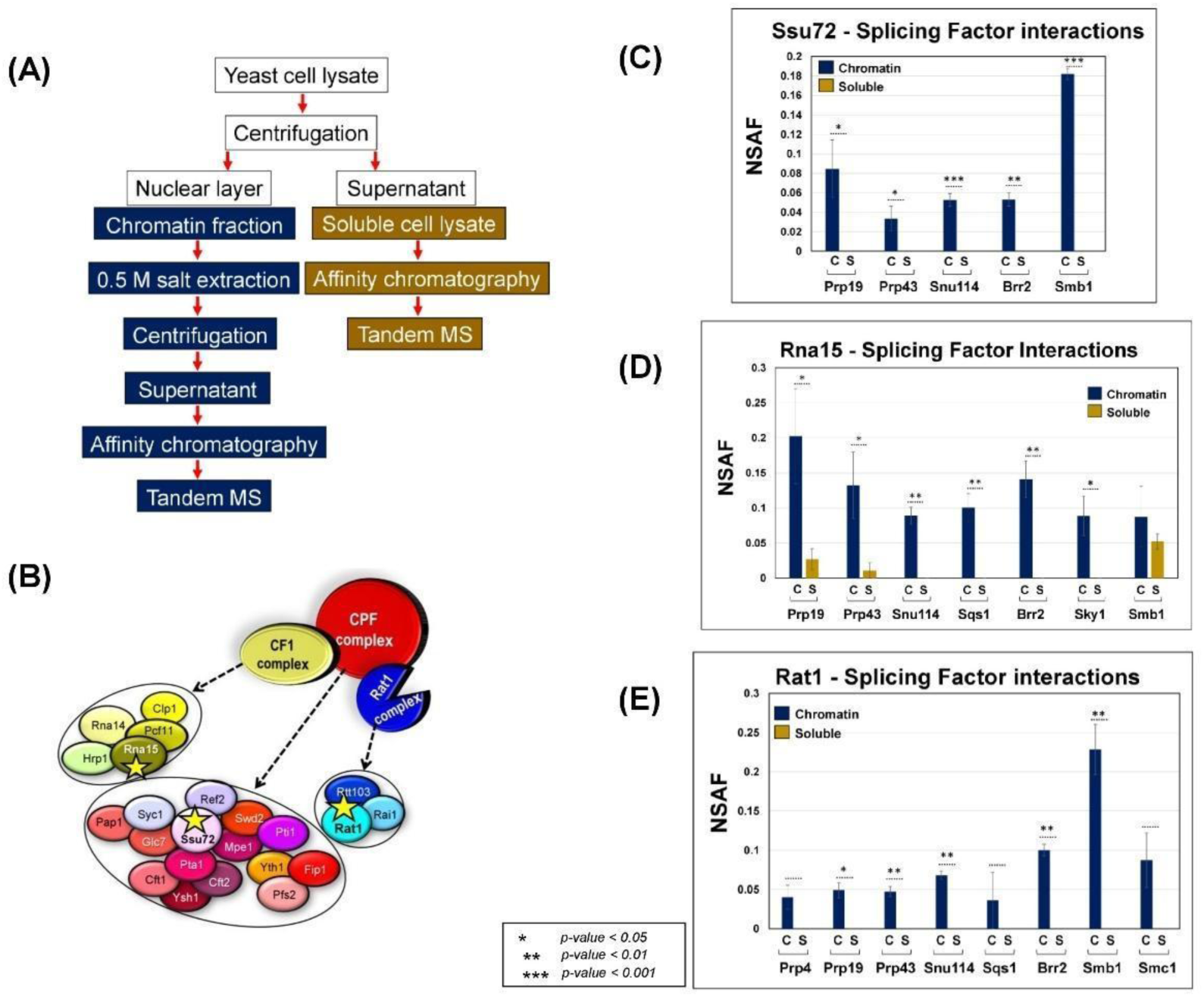
Purified termination complexes interact with splicing factors exclusively in the chromatin fraction. **(A)** The workflow for identifying termination factor-associated proteins in affinity purified preparations derived from chromatin and soluble fractions. **(B)** Schematic representation of CPF, CF1 and Rat1 termination complexes of yeast showing their respective subunits. The subunit that was epitope-tagged for affinity purification is marked with an asterisk. **(C)** Splicing factors interacting with the Ssu72 subunit of CPF complex. **(D)** Splicing factors interacting with the Rna15 subunit of CF1 complex. **(E)** Splicing factors interacting with the Rat1 subunit of Rat1complex. All splicing factor-termination factor interactions were observed in the chromatin context (blue bar). *p*-values calculated by two tailed t-test indicate significant enrichment of the splicing factors factor in chromatin fraction relative to soluble fraction. Error bars represent one unit of standard deviation based on four biological replicates.

### CPF, CFI and Rat1 termination complexes interact with RNAPII and the general transcription factors

It is known that all three termination complexes are recruited to chromatin cotranscriptionally as RNAPII reaches the 3’ end of a gene. We, therefore, assessed the presence of RNAPII subunits in the affinity purified termination complex preparations. All three termination complexes exhibited interaction with multiple RNAPII subunits (Supplemental Fig. 1). Notably, these interactions were observed exclusively in the chromatin derived preparations of termination factors, thereby emphasizing their transcription-dependent nature (Supplemental Fig. 1). We next investigated interactions of termination complexes with the general transcription factors. The results corroborated the previously published interaction of CF1 and Rat1 complexes with TFIIB (Supplemental Fig. 2B and 2C). Additionally, all three termination complexes interact with subunits of the TFIID complex (Supplemental Fig. 2A, 2B and 2C). Combined these findings presented unambiguous evidence of a novel role of TFIID in promoter-terminator crosstalk, with TFIID engaging in stronger interactions with all three termination complexes compared to TFIIB. These results corroborated the preliminary evidence from earlier studies that there is extensive crosstalk between promoter and terminator-bound factors during transcription, which may be crucial for successful accomplishment of RNAPII transcription cycle and cotranscriptional RNA processing.

### Termination complexes also interact with splicing factors

In yeast, splicing often occurs rapidly within seconds following the transcription of an intron, while the polymerase is just a few nucleotides downstream of the 3′ splice site (Oesterreich et al., 2016; Herzel et al., 2017; Neugebauer, 2019). Given the cotranscriptional nature of splicing, we reasoned that factors involved in initiation, elongation, and termination of transcription may physically or functionally interact with splicing factors. Consistent with this idea, we recently demonstrated interactions between the general transcription factor TFIIB and the termination factor Rat1 with several splicing factors in the affinity-purified preparations (Dhoondia et al., 2021; O’Brien and Ansari, 2024). We therefore investigated the presence of splicing factors in our purified termination factors preparations from both chromatin and soluble cellular fractions. These efforts uncovered evidence that all three 3’ end processing-termination complexes interact with splicing factors (Fig. 1C, 1D and 1E). Most of these interactions were observed exclusively in the chromatin context, indicating their transcription-dependent nature (Fig. 1C, 1D and 1E dark blue bars). Notably, the NSAF values for these interactions ranged from 0.05 to 0.2, suggesting that splicing factors were present in sub-stoichiometric amounts in the termination factor preparations.

Despite being stable at 0.5 M ammonium sulfate, these observed interactions are transient. It is particularly notable that all three termination complexes interacted with a similar set of splicing factors, including Prp19, Prp43, Snu114, Brr2, and Smb1 (Fig. 1C, 1D and 1E). In addition, Ssu72 also interacted with Smd2 (Fig. 1C) and Rna15 with Sqs1, Sky1 and Smb1 (Fig. 1D), while Rat1 made additional contacts with Prp4 and Sqs1 (Fig. 1E). With the exception of a few non-snRNP proteins, most of these splicing factors are components of snRNP complexes. This included Prp19, which is a component of the NineTeen complex (NTC), which functions during catalytic activation of the spliceosome (Chan et al., 2003; Chan and Cheng, 2005; Hogg et al., 2010) and Prp43, an RNA helicase, which is involved in removing U2, U5, and U6 snRNPs from the post-splicing lariat-intron ribonucleoprotein complex (Bohnsack et al., 2022).

### Splicing factor Prp19 interacts with multiple factors linked to transcription

To explore the significance of the Prp19 and Prp43 interactions in transcription and cotranscriptional RNA processing in more detail, we performed reciprocal purification experiments. We epitope-tagged Prp19 and Prp43 and performed affinity purification from chromatin and soluble cellular fractions. Mass spectrometric analysis of these preparations produced both expected and surprising results. While no significant interaction with the transcription machinery was detected for Prp43 (Supplemental Fig. 3), Prp19 showed robust, statistically significant interactions with RNAPII (Fig. 2A), TFIID (Fig. 2B) and the CPF termination complex (Fig. 2D). A related unexpected finding was evidence for a strong interaction of Prp19 with the RSC chromatin remodeling complex (Fig. 2C).

**Figure 2:**
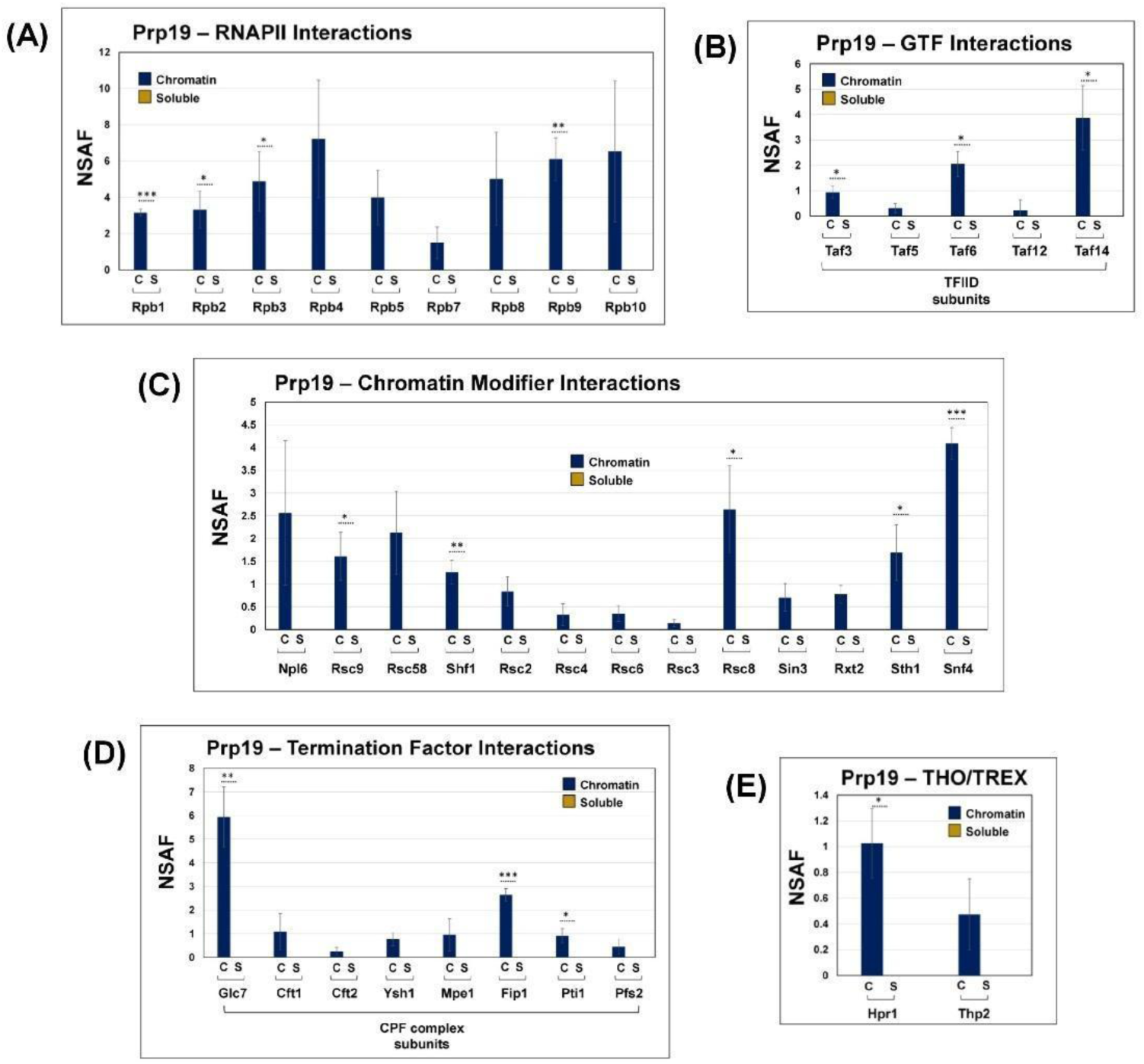
Prp19 interacts with RNAPII, general transcription factor TFIID, RSC chromatin modifiers and components of the TREX complex in chromatin fraction. **(A)** Prp19 exhibits strong interaction with nine of the twelve subunits of RNAPII in the chromatin fraction. **(B)** Prp19 associates with four subunits of the general transcription factor TFIID. **(C)** Prp19 interacts with almost entire RSC chromatin remodeling complex. **(D)** Prp19 interacts with multiple subunits of the CPF termination complex. **(E)** Prp19 interacts with two subunits of the TREX complex. *p*-values calculated by two tailed t-test indicate significant enrichment of the splicing factors factor in chromatin fraction relative to soluble fraction. One asterisk (*) signifies a p-value equal to or smaller than 0.05 (p ≤ 0.05); two asterisks (**) signify a p-value equal to or smaller than 0.01 (p ≤ 0.01); while three asterisks (***) signify a p-value equal to or smaller than 0.01 (p ≤ 0.001). Error bars represent one unit of standard deviation based on four biological replicates.

The splicing factors are cotranscriptionally recruited by the CTD of RNAPII (Misteli and Spector, 1999; Nojima et al., 2018; Guo et al., 2019). There are also reports of interaction between components of the Prp19 containing NTC with RNAPII (Chanarat et al., 2011). We therefore examined the interaction of Prp19 with RNAPII subunits. Our results sshowed strong, stable interaction of Prp19 with at least nine subunits of RNAPII: Rpb1, Rpb2, Rpb3, Rpb4, Rpb5, Rpb7, Rpb8, Rpb9 and Rpb10 (Fig. 2A). The high NSAF values indicate that the interaction of Prp19 with RNAPII subunits is stronger than that of any of the termination factors described above. Next, we assessed the representation of GTFs in the purified Prp19 preparations. Although several GTFs including TFIIB were detected, only TFIID subunits, particularly Taf3, Taf5, Taf6, Taf12 and Taf14, exhibited a statistically significant enrichment in chromatin derived preparation (Fig. 2B). The TFIID interaction may have implications in both transcription and splicing. We therefore also probed for the presence of termination factors in the Prp19 preparation. Although Prp19 was detected in the chromatin derived preparations of CF1, CPF and Rat1 termination complexes, only CPF subunits were recorded with a stable, statistically significant presence in the affinity purified chromatin Prp19 preparation (Fig. 2D). Moreover, the CPF subunits Glc7, Cft1, Cft2, Ysh1, Mpe1, Fip1, Pti1 and Pfs2 were consistently detected in a stoichiometric amount (Fig. 2D).

The most notable Prp19 interaction, however, was observed with RSC chromatin remodeling complex. Prp19 exhibited robust interaction with almost all subunits of the RSC complex. The RSC subunits Sth1, RSC2, RSC3, RSC4, RSC6, RSC8, RSC9, RSC58 were highly enriched in the chromatin-derived Prp19 preparation (Fig. 2C). RSC complex is especially critical for initiation of transcription by virtue of its role in remodeling +1 nucleosome (Lorch and Kornberg, 2017; Neumann and Wilkins, 2021). The complex has also been implicated in elongation of transcription (Biernat et al., 2021). These interactions were observed exclusively in the chromatin fraction, underscoring the transcription-dependent nature of the interaction. Taken together, these findings suggested that Prp19 may play a previously unreported role in the initiation and termination step of transcription.

### Evidence of Prp19 interaction with the RNA export complex TREX

Prp19 is a part of NTC (Chanarat and Sträßer, 2013; Idrissou and Marechal, 2022), which is an evolutionarily conserved complex with homologs reported form both yeast and humans. It was discovered as a complex that functions in splicing during the catalytic activation of the spliceosome (Cheng et al., 1993; Tarn et al., 1994; Chan et al., 2003; Chan and Cheng, 2005; Hogg et al., 2010). In a previous study, mass spectrometry of affinity purified NTC/Prp19C from the spliceosomal fraction of yeast led to the identification of nine subunits: Prp19, Cef1, Snt309, Prp46, Syf1, Syf2, Syf3/Clf1, Isy1 and Ntc20 (Ohi and Gould, 2002; Fabrizio et al., 2009). Furthermore, the Prp19 complex also interacts with the THO complex in yeast, which is a subcomplex of TREX, a conserved complex that couples transcription to nuclear mRNA export (Jimeno et al., 2002; Chanarat et al., 2011; Henke-Schulz et al., 2024). We therefore examined the presence of NTC and TREX complex subunits in the affinity purified Prp19 preparations. While we failed to detect statistically significant evidence of the presence of any of the NTC/Prp19C subunits in either soluble or the chromatin derived preparations, two subunits of the TREX complex, Hpr1 and Thp2, were consistently detected with high confidence in the chromatin derived Prp19 preparation (Fig. 2E). These results suggested that Prp19 may interact with multiple protein complexes depending on its functional state. Although the NTC subunits Syf1 and Syf2 have been implicated in elongation of transcription, no conclusive evidence has been previously reported regarding a direct role of Prp19 in either elongation or any other step of the transcription cycle (Chanarat et al., 2011; Henke-Schulz et al., 2024). The interaction of Prp19 with RNAPII, TFIID, CPF complex as well as the RSC complex therefore indicated a novel role of Prp19 in the initiation and termination steps of transcription.

### Prp19 interaction with RNAPII, TFIID and TREX complex is not mediated by DNA or RNA

Our findings that Prp19 interacts with RNAPII, TFIID, the CPF complex, the TREX complex, and the RSC complex specifically in the chromatin context led us to speculate that these interactions might be indirect, potentially mediated by the template DNA or the transcribing mRNA. To investigate whether these interactions were dependent on nucleic acids, we subjected chromatin eluate to micrococcal nuclease (MNase) digestion before performing affinity chromatography and mass spectrometry (O’Brien and Ansari, 2024). MNase digests chromatin by cleaving exposed DNA between nucleosomes and RNA, thereby releasing any protein-DNA or protein-RNA complexes.

After MNase digestion, we observed that the interaction between Prp19 and most of the subunits of RNAPII, TFIID, and the TREX complex remained largely unaffected (Table 1A, 1B, 1D and 1E). This suggested that Prp19 directly associates with these complexes independent of any intermediary nucleic acids. Notably, however, the interaction with specific subunits, such as Rpb10 of RNAPII and Taf3 of TFIID, was completely abolished (Table 1A, 1B). Additionally, interactions with Taf14 and Hpr1 showed a marked decrease (Table 1B and 1D). These observations indicated that Prp19’s association with these complexes was not generally dependent on DNA or RNA, but that certain subunits may still be partially reliant on nucleic acids for maintaining a stable interaction. In contrast, Prp19 interaction with several subunits of the CPF and RSC complexes was significantly reduced or completely lost after MNase treatment (Table 1C, 1E). This suggested that the binding of Prp19 to these complexes was, to some extent, dependent on nucleic acids, either directly or by stabilizing protein-protein interactions.

**Table 1:** The table shows NSAF values for all interacting partners of Prp19 shown in Fig. 2, before and after MNase digestion. The red boxes indicate that the interaction of Prp19 with the protein was completely abolished in the presence of MNase, while yellow boxes indicate that interaction of the protein with Prp19 was compromised in the presence of MNase. White boxes depict the interaction being completely unaffected by MNase.

| (A) |  |  |
| --- | --- | --- |
| RNAPII subunits | Chromatin +MNase | Chromatin -MNase |
| Rpb1 | 7.17±0.61 | 3.17±0.18 |
| Rpb2 | 2.26±0.31 | 3.31±1.01 |
| Rpb3 | 2.17±1.44 | 4.89±1.63 |
| Rpb4 | 9.88±1.52 | 7.22±3.25 |
| Rpb5 | 9.12±1.31 | 3.99±1.51 |
| Rpb7 | 3.00±1.73 | 1.50±0.86 |
| Rpb8 | 7.82±1.12 | 5.03±2.56 |
| Rpb9 | 6.11±1.17 | 6.11±1.17 |
| Rpb10 | 0 | 6.53±3.89 |

| (B) |  |  |
| --- | --- | --- |
| GTFs | Chromatin +MNase | Chromatin -MNase |
| Taf3 | 0 | 0.95±0.24 |
| Taf5 | 1.39±0.28 | 0.32±0.18 |
| Taf6 | 2.95±0.57 | 2.05±0.50 |
| Taf12 | 2.65±1.33 | 0.23±0.40 |
| Taf14 | 1.76±0.35 | 3.87±1.26 |

| (C) |  |  |
| --- | --- | --- |
| Termination factors | Chromatin +MNase | Chromatin -MNase |
| CPF complex |  |  |
| Glc7 | 8.97±0.53 | 5.94±1.27 |
| Cft1 | 0 | 1.09±0.76 |
| Cft2 | 0.20±0.19 | 0.25±0.17 |
| Ysh1 | 0 | 0.76±0.29 |
| Mpe1 | 0 | 0.95±0.68 |
| Fip1 | 3.17±0 | 2.64±0.26 |
| Pti1 | 0 | 0.91±0.30 |
| Pfs2 | 0 | 0.45±0.32 |

| (D) |  |  |
| --- | --- | --- |
| THO and TREX subunits | Chromatin +MNase | Chromatin -MNase |
| Hpr1 | 0.43±0.21 | 1.03±0.26 |
| Thp2 | 2.53±0.63 | 0.47±0.27 |
| Sub2 | 14.25±2.96 | 0.57±2.57 |

| (E) |  |  |
| --- | --- | --- |
| Chromatin Modifiers | Chromatin +MNase | Chromatin -MNase |
| Npl6 | 1.14±0 | 2.56±1.58 |
| Rsc9 | 0.58±0.29 | 1.61±0.53 |
| Rsc58 | 0.66±0.33 | 2.13±0.91 |
| Shf1 | 0 | 1.26±0.26 |
| Rsc2 | 0 | 0.84±0.32 |
| Rsc4 | 0.26±0.26 | 0.33±0.23 |
| Rsc6 | 0.18±0.17 | 0.35±0.17 |
| Rsc3 | 0 | 0.14±0.08 |
| Rsc8 | 2.26±0.69 | 2.64±0.96 |
| Sin3 | 1.30±0.19 | 0.71±0.30 |
| Rxt2 | 2.33±0.67 | 0.78±0.19 |
| Sth1 | 0 | 1.69±0.61 |
| Snf4 | 1.32±0.26 | 4.09±0.35 |

To further validate the specificity of these interactions, we assessed the presence of histones in the purified Prp19-chromatin preparation. The lack of statistically significant interactions of histones with Prp19 strongly supported the conclusion that Prp19 is not indiscriminately interacting with proteins bound to chromatin (Supplemental Fig. 4) and that its interactions are highly selective and specific instead, reinforcing the authenticity of the identified protein-protein associations.

### Prp19 has a novel role in transcription by RNAPII

The strong interaction of Prp19 with TFIID and RSC complex in the chromatin context gave rise to the speculation that Prp19 may have a role in transcription by interacting with RNAPII. While Syf1 and Syf2 subunits of Prp19C have been implicated in RNAPII transcription cycle at the elongation step (Chanarat et al., 2011; Henke-Schulz et al., 2024), the direct involvement of Prp19 itself in the transcription cycle has not been demonstrated so far. To address this possible gap, we performed auxin-mediated depletion of Prp19 in yeast cells as described in Shetty et al., (2019). The auxin-inducible degron (AID) is a powerful tool for depletion of essential proteins to study their function *in vivo* in non-plant eukaryotes. This method can conditionally induce the degradation of any protein by the proteasome, simply by the addition of the plant hormone auxin. Application of the AID protocol resulted in almost complete degradation of Prp19 within 90 minutes of adding auxin to the medium (Fig. 3A). Furthermore, the growth of Prp19-AID tagged yeast cells was adversely affected in the presence of auxin in the medium (Fig. 3B). No such growth defect was observed for the isogenic wild-type strain under the same conditions (Fig. 3B).

**Figure 3:**
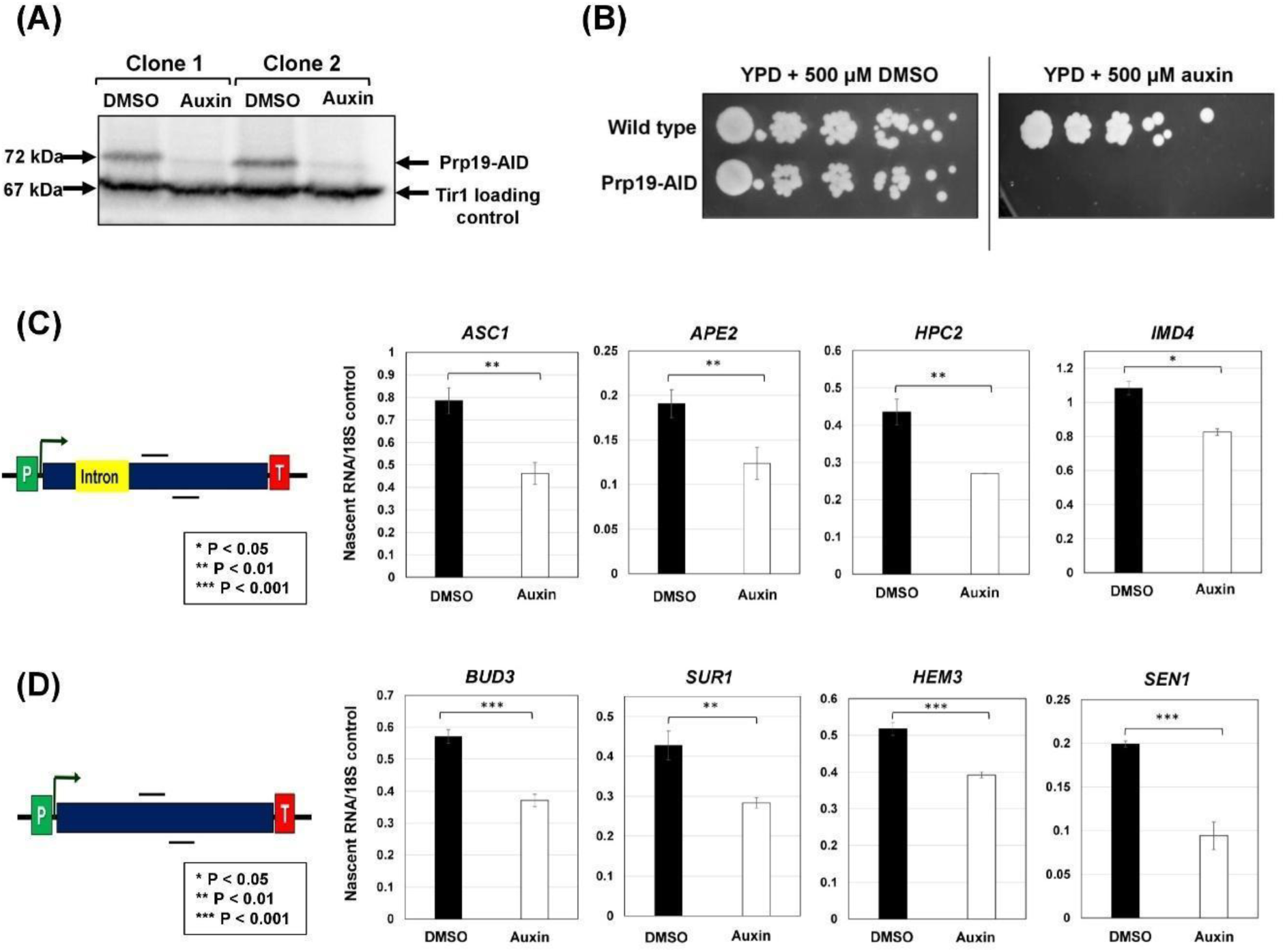
Prp19 affects transcription of both intron-containing and intron-lacking genes. **(A)** Western blot showing auxin-mediated depletion of AID-tagged Prp19 from yeast cells. **(B)** Serial dilution plating assay showing complete loss of cell viability in the presence of 500 mM of auxin. **(C)** Nascent mRNA level of intron-containing genes; *ASC1*, *APE2*, *HPC2*, and *IMD4,* exhibits a 15-40% decline upon auxin-mediated depletion of Prp19. (D) Nascent RNA level of intron-lacking genes; *BUD3*, *SUR1*, *HEM3*, and *SEN1* exhibited a 2-50% reduction in the absence of Prp19. The nascent transcript level of 18S rRNA was used as the normalization control. *p*-values calculated by two tailed t-test indicate significant enrichment of the splicing factors factor in chromatin fraction relative to soluble fraction. Error bars represent one unit of standard deviation based on four biological replicates.

To examine the role of Prp19 in transcription, we performed Transcription Run-On (TRO) assay in Prp19-AID-tagged strains after 90 minutes of Prp19 depletion in the presence of auxin. We monitored transcription of four intron-containing genes (*ASC1, APE2, HPC2* and *IMD4*) and four intron-lacking genes (*BUD3, SUR1, HEM3* and *SEN1*). The transcription of the four intron-containing genes decreased by about 25-40% (Fig. 3C), while that of four intron-lacking genes registered a 25-50% decline upon auxin-mediated depletion of Prp19 (Fig. 3D). These results provided strong evidence that Prp19 affects transcription of at least a subset of both intron-containing and intron-lacking genes in yeast.

### Genome-wide evidence of an intron-independent role of Prp19 in gene transcription

Next, we investigated Prp19’s involvement in transcription on a genome-wide scale. To achieve this, we employed the ‘Global Run-On-Seq’ (GRO-seq) approach, which is a genome-wide, strand-specific adaptation of the traditional transcript run-on (TRO) method (Core et al., 2008), The method provides a high-resolution snapshot of transcriptional activity, allowing for precise mapping of the position and density of actively transcribing RNA polymerase in a strand-specific manner. GRO-seq was performed in both wild-type and Prp19-depleted yeast cells, with three biological replicates per condition. The sequencing reads were aligned to the S. cerevisiae S288c genome (version R64-1-1) obtained from SGD. To focus our analysis on transcriptionally active genes under standard YPD growth conditions, we selected mRNAs with mean expression levels of ≥10 reads.

To investigate the differential expression of genes in wild-type and Prp19-depleted cells, we generated a mean-difference (MA) plot (Fig. 4A). This plot displays the relationship between the log fold-change and the mean expression levels in both the wild-type and Prp19-depleted conditions. The Y-axis represents the base-2 log fold-change, while the X-axis shows the normalized mean expression. Data points that are located at the extremes along the Y-axis indicate genes with highly differential expression levels. For this analysis, we used genes that were separated from their neighboring genes by at least 250 nucleotides. This was done to remove the interfering effect of neighboring genes in our analysis. Out of these 3140 genes, 2918 had mean expression level greater than 10 and were used for further analysis. Genes exhibiting a 2-fold or greater change in expression upon Prp19 depletion are highlighted in red, while those with less than a 2-fold change, or no significant change, are shown in grey. This approach indicated that nearly 193 genes were upregulated, and 217 genes were downregulated following the depletion of Prp19 (Fig. 4A). However, a considerably larger group of genes showed more subtle changes in expression, ranging from 25-50%, which is less than a 2-fold change (Fig. 4A), pointing at a possibly broader effect of Prp19 on the global gene expression landscape of yeast.

**Fig. 4.**
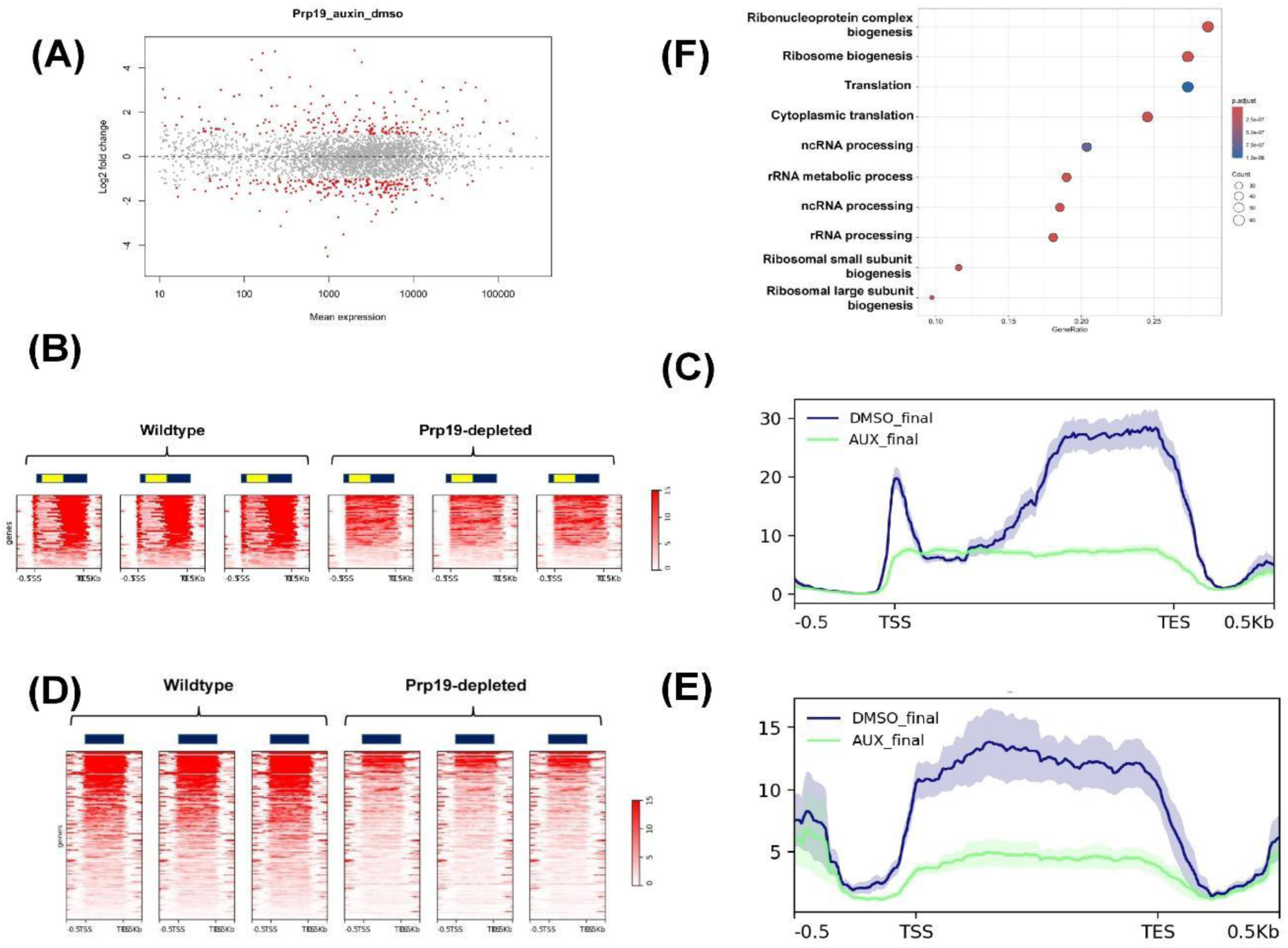
GRO-Seq demonstrates genomewide alteration in transcription upon auxin-mediated depletion of Prp19 from yeast cells. **(A)** MA plot displaying the relationship between the log_2_ fold-change and the mean expression levels of genes derived from GRO-Seq reads in wild-type (WT) and Prp19-depleted cells. Genes that are significantly upregulated and downregulated are highlighted in red. The Y-axis represents the base-2 log fold-change, while the X-axis shows the normalized mean expression. Data represents the average of three biological replicates for each condition. **(B)** Heat maps show GRO-seq signal of the 93 downregulated intron-containing genes. The read densities are from −500 nucleotides upstream of TSS and +500 nucleotides downstream of the TES in WT and Prp19-depleted cells. All three replicates are shown here. The genes are arranged from top to bottom in descending order of relative expression. **(C)** Metagene plot of normalized read densities of intron-containing genes derived from heat maps in ‘B’ in WT and Prp19-depleted cells. Thick/center lines are the average normalized read densities. Upper and lower light-colored lines represent standard error. **(D)** Heat maps show GRO-seq signal of the 302 downregulated intron-lacking genes from −500 nucleotides upstream of TSS and +500 nucleotides downstream of the TES in WT and Prp19-depleted cells in three biological replicates. **(E)** Metagene plot of normalized GRO-Seq read densities of intron-lacking genes derived from heat maps in ‘D’ in WT and Prp19-depleted cells. **(F)** Ontological analysis of genes downregulated upon Prp19 depletion from yeast cells.

In the next step of our analysis, we focused on genes that required Prp19 for reaching optimal expression levels, suggested by their downregulation in Prp19 depleted cells. Given that Prp19 is a splicing factor, and that splicing effects transcription of intron-containing genes, we categorized downregulated genes into ‘intron-containing’ and ‘intron-lacking’ groups. Of the 217 genes downregulated in the absence of Prp19, 53 were intron-containing, while 164 are intron-lacking, corroborating the evidence that Prp19 regulates transcription in a manner that is independent of its role in splicing.

The heatmap of the 53 significantly downregulated intron-containing genes revealed a consistent reduction in transcription across the exonic regions in all three replicates in the absence of Prp19 (Fig. 4B). Moreover, metaplot analysis indicated a 2-3-fold decrease in transcription of the same loci upon Prp19 depletion (Fig. 4C). Notably, heatmaps of the 164 intron-lacking genes exhibited a similar reduction in transcription across coding regions in the absence of Prp19 (Fig. 4D) and the metaplot for these genes further confirmed a three-fold or greater decrease in transcription following Prp19 depletion (Fig. 4E).

These results collectively suggested that Prp19 is essential for the optimal transcription of a subset of both intron-containing and intron-lacking genes. However, it is important to note that Prp19 is not a general transcription factor like those traditionally involved in basal transcription initiation. This distinction suggests that Prp19 might play a more specialized or context-dependent role in transcription regulation.

### Differential gene expression evidence of a specific role of Prp19 in cell growth regulation

To explore the shared features of genes dependent on Prp19 for transcription, we searched for common structural or sequence motifs. Interestingly, however, genes whose transcription was reduced upon Prp19 depletion did not exhibit a clear pattern in terms of predicted structures or enriched sequence motifs. Additionally, Prp19-associated transcriptional reduction was not linked to the presence or absence of a TATA box in the promoter region. Both TATA-box-containing and TATA-box lacking genes exhibited Prp19-dependence, suggesting that Prp19’s role in transcription is not confined to a particular type of promoter architecture.

Ontological analysis of Prp19-dependent genes revealed that transcription of genes involved in translation, ribosomal subunit biogenesis, and ribosome assembly were particularly sensitive to Prp19 depletion (Fig. 4F), indicating a pivotal role of the protein in maintaining the transcription of genes that are essential for protein synthesis and cellular growth. In contrast, ontological analysis of genes with enhanced transcription in the absence of Prp19 revealed a strong association with processes related to cell wall and membrane biology (Supplemental Fig. 5). This suggested that Prp19 may exert an inhibitory effect on the transcription of gene sets involved in cell wall and membrane-related processes, further consistent with a role in fine-tuning transcriptional responses under different cellular conditions.

Overall, these findings provided compelling evidence that Prp19 is not only a splicing factor but also plays a critical, albeit specialized, role in transcription regulation. The ability of Prp19 to influence the transcription of both intron-containing and intron-lacking genes opens new avenues for understanding the complex interplay between splicing and transcription. The association between Prp19 and processes like ribosome biogenesis further highlights its potential role in coordinating the transcriptional and translational machinery necessary for cell growth and function. This dual functionality of Prp19 underscores the importance of moonlighting factors in cellular processes and highlights a novel aspect of transcriptional regulation that warrants further exploration.

### Prp19 affects the initiation step of transcription

To determine which step or steps of transcription were affected by Prp19, we focused on its interaction with TFIID and the RSC complex, which implied functions transcription initiation. To this end, we monitored the recruitment of the TFIID complex to the promoter regions of three intron-containing and three intron-lacking genes in the presence and absence of Prp19. Recruitment was assessed by chromatin immunoprecipitation (ChIP) using an antibody against the TBP subunit of TFIID and PRC reaction using primer pairs shown in Figs 5A and 5C. TBP occupancy at the promoters of both gene categories decreased by approximately 65-80% in the absence of Prp19 (Fig. 5B and 5D). Although transcription of these genes was reduced by just 25-50% in the absence of Prp19, a substantial decrease in TBP recruitment upon Prp19 depletion was observed. As we expected that genes whose transcription is affected by two-fold or more in the absence of Prp19 will exhibit a more pronounced initiation defect, these results provided strong evidence that Prp19 plays a novel role in the initiation of transcription. However, it remains possible that Prp19 may also be involved in the elongation step of transcription, though this cannot be definitively concluded from these data.

**Figure 5:**
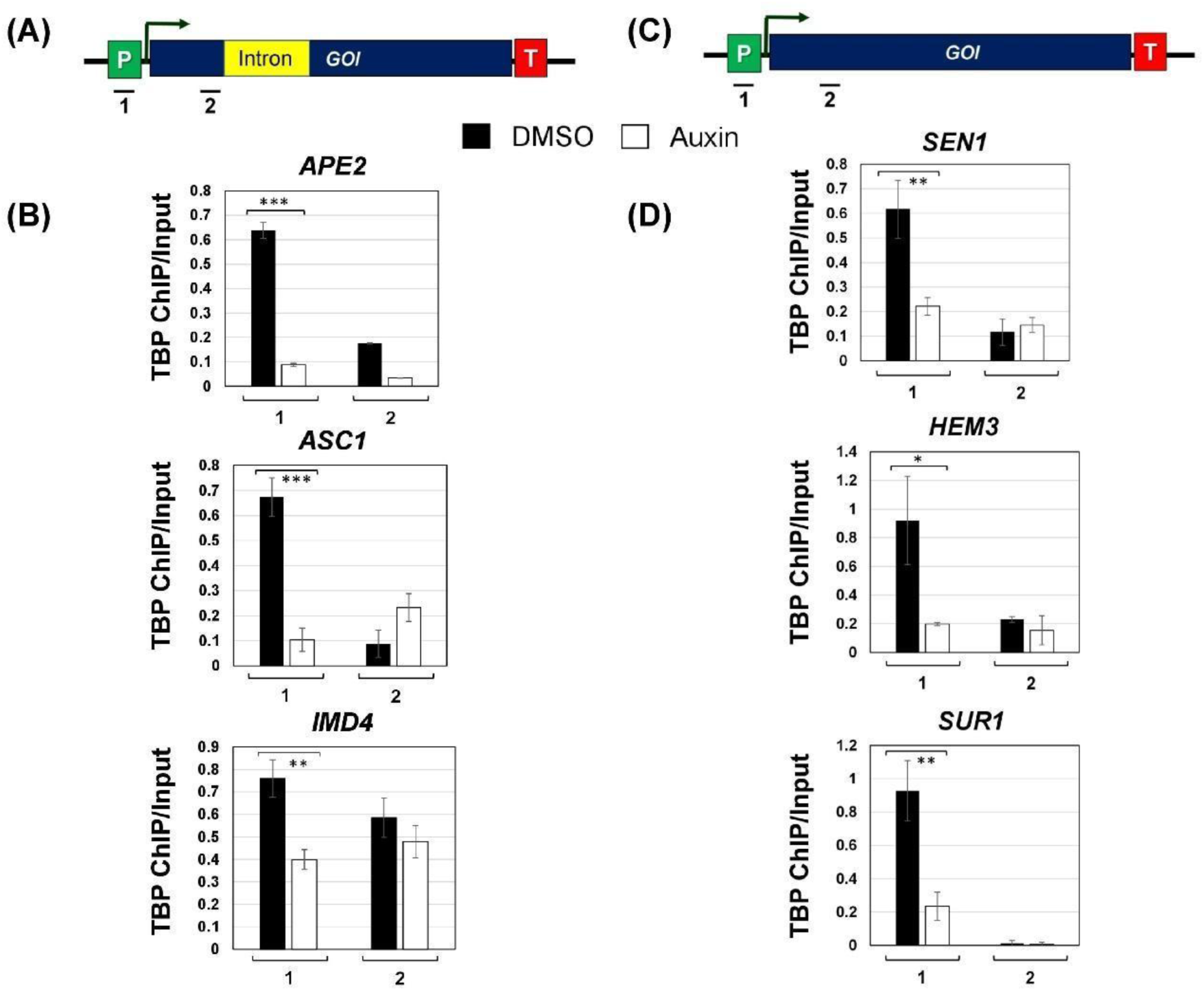
Promoter occupancy of TATA-binding protein (TBP) is adversely affected in the absence of Prp19. **(A)** Schematic depiction of an intron-containing gene showing the position of the primer pairs 1 and 2 used in ChIP assay. **(B)** The intron-containing genes; *APE2*, *ASC1*, and *IMD4* display a statistically significant decrease in promoter occupancy of TBP upon auxin-mediated depletion of Prp19 from cells. **(C)** Schematic depiction of an intron-lacking gene showing the position of the primer pairs 1 and 2 used in ChIP assay. **(D)** The intron-lacking genes; *SEN1*, *HEM3*, and *SUR1* exhibit a decreased occupancy of TBP at the promoter region in the absence of Prp19. The Input signal, representing DNA prior to immunoprecipitation, was used as normalization control. ChIP results presented here represent the average of three biological and five technical replicates. p-values calculated by two tailed t-test indicate significant enrichment of the splicing factors factor in chromatin fraction relative to soluble fraction. One asterisk (*) signifies a p-value equal to or smaller than 0.05 (p ≤ 0.05); two asterisks (**) signify a p-value equal to or smaller than 0.01 (p ≤ 0.01); while three asterisks (***) signify a p-value equal to or smaller than 0.01 (p ≤ 0.001). Error bars represent one unit of standard deviation based on four biological replicates.

Prp19 may be playing a direct role in transcription of RNAPII-transcribed genes, or it may be affecting transcription indirectly. To distinguish between these possibilities, we performed Prp19-ChIP for three intron-containing and three intron-lacking genes, which showed reduced transcription upon auxin-mediated depletion of Prp19. We reasoned that if Prp19 is directly involved in transcription initiation, it would be recruited to the promoter region of these genes. The ChIP-PCR was performed using primer pairs shown in Figs 6A and 6C. The ChIP results revealed that Prp19 crosslinked to the promoter, coding region as well as the terminator region of all six genes under investigation (Fig. 6B and 6D). Notably, there was a peak of Prp19 at the promoter region, as well as a minor peak at the terminator region of all six genes (Fig. 6B and 6D). Together, these findings corroborated that major conclusion that Prp19 plays a direct role in the transcription of both intron-containing and intron-lacking genes in yeast.

**Figure 6:**
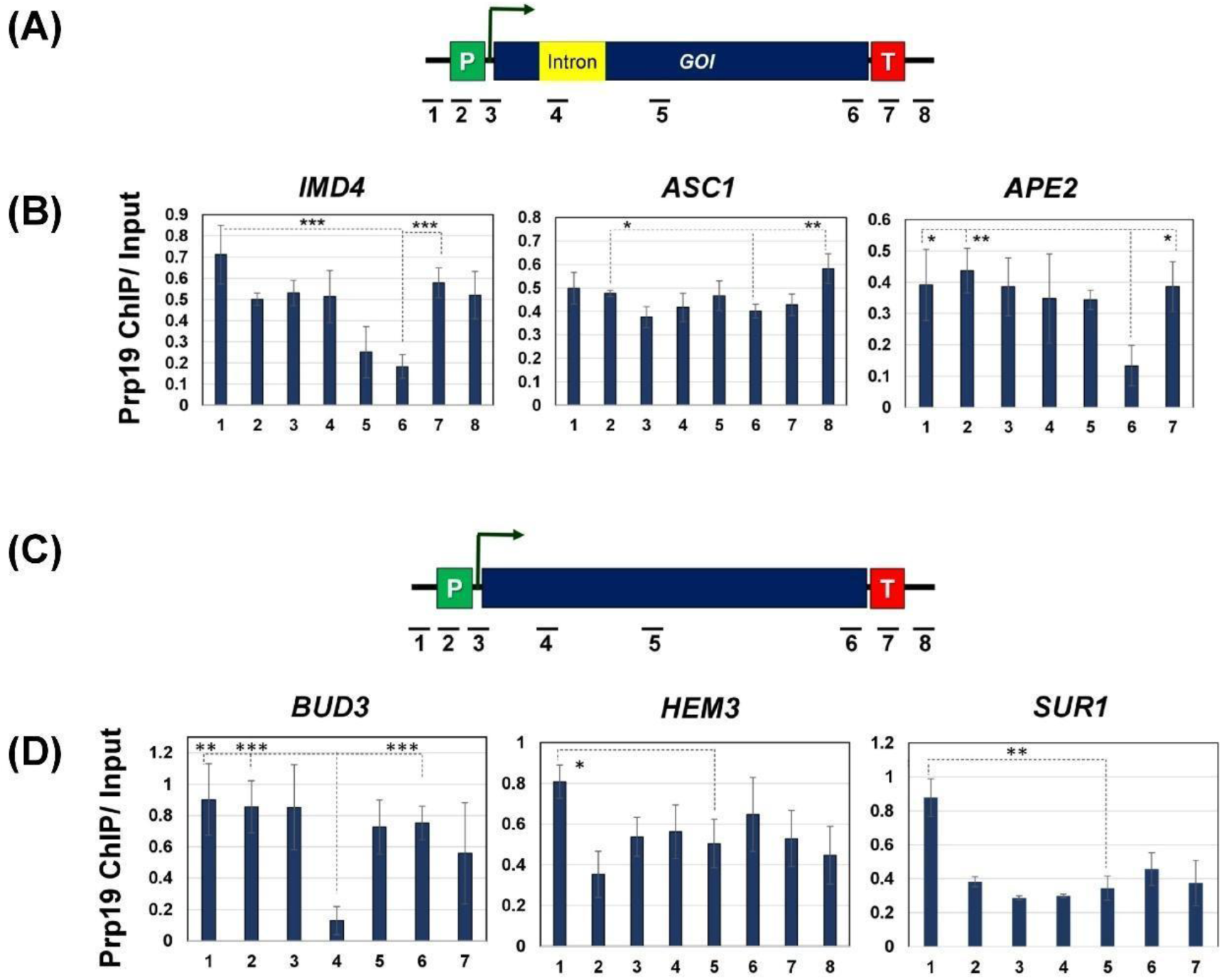
Prp19 is recruited to the promoter, terminator and coding region of both intron-containing and intron-lacking genes. **(A)** Schematic depiction of an intron-containing gene showing the position of the primer pairs used in ChIP assay. **(B)** Intron-containing genes *IMD4*, *ASC1*, and *APE2* show a moderate increase in Prp19 occupancy at the promoter and terminator regions compared to the body of the gene. **(C)** Schematic depiction of an intron-lacking gene showing the position of the primer pairs used in ChIP assay. **(D)** The intron-lacking genes *BUD3*, *HEM3*, and *SUR1* show an increase in Prp19 occupancy at the promoter region as compared to the body of the gene. The Input signal, representing DNA prior to immunoprecipitation, was used as normalization control. ChIP results presented here represent the average of three biological and five technical replicates. p-values calculated by two tailed t-test indicate significant enrichment of the splicing factors factor in chromatin fraction relative to soluble fraction. One asterisk (*) signifies a p-value equal to or smaller than 0.05 (p ≤ 0.05); two asterisks (**) signify a p-value equal to or smaller than 0.01 (p ≤ 0.01); while three asterisks (***) signify a p-value equal to or smaller than 0.01 (p ≤ 0.001). Error bars represent one unit of standard deviation based on four biological replicates.

## Discussion

The classical view of transcription and cotranscriptional RNA processing is that factors involved in these processes have dedicated step-specific functions. Transcription initiation and termination factors are seen as key to the beginning and end of RNA synthesis, while splicing factors are traditionally confined to their role in intron removal during RNA maturation. A number of studies, however, have increasingly challenged this dogma, revealing that many of these factors participate in broader networks that integrate transcription and RNA processing. For instance, general transcription factor TFIIB, traditionally associated with transcription initiation, is now known to play a role in 3’ end processing/transcription termination (O’Brien et al., 2024; Santana et al., 2022). Likewise, factors implicated in 3’ end processing, including the yeast CF1 complex and mammalian CPSF and CstF, have been shown to affect promoter-associated transcription (Tan-Wong et al., 2012; Al-Husini et al., 2013; Almada et al., 2013; Ntini et al., 2013). A growing body of evidence suggests that transcription and RNA processing are not independent, temporally separate events but are intricately coupled through a network of interactions among a variety of accessory proteins including RNA-binding proteins (Robert et al., 2002; Kornblihtt et al., 2004; Bentley, 2014; Custodio and Carmo-Fonseca, 2016; Herzel et al, 2017; Neugebauer, 2019; Saidi et al., 2016; Tellier et al., 2020; Wallace and Beggs 2017; Shenasa and Bentley, 2023). Notably, splicing factors have been shown to influence transcription, and transcription factors have been found to modulate splicing, blurring the boundaries between these processes (Fong and Zhou, 2001; Rambout et al., 2018; Shao et al., 2020; Leader et al., 2021; Caizzi et al., 2021; Carnesecchi et al., 2022; Göös et al., 2022; Oksuz et al., 2023; Chanarat et al., 2011; Henke-Shulz et al., 2024). Here, we present data that further integrate transcription and splicing, showing that the splicing factor Prp19, a key player in spliceosome assembly, also plays a novel and direct role in transcription initiation. Furthermore, Prp19 affected transcription initiation of both intron-containing and intron-lacking genes, thereby underscoring the splicing-independent role of the factor in transcription. The possibility of Prp19 being a transcription factor with additional function in splicing cannot be ruled out. Our findings provide new insight into how splicing factors, traditionally considered separate from transcription, contribute to the regulation of gene expression from the very start of the transcription cycle.

Prp19 complex (NTC/Prp19C) is a multiprotein complex consisting of nine core proteins and up to 19 associated proteins in yeast (Ohi et al., 2002; Fabrizio et al., 2009). In humans, more than 30 proteins have been identified in the homologous Prp19C complex (Cheng et al., 1993; Tarn et al., 1994; Chan et al., 2003; Chan and Cheng, 2005; Hogg et al., 2010). The components of yeast NTC/Prp19C were identified using a combination of genetic and biochemical approaches. The biochemical approach involved isolation of active yeast spliceosomes by centrifugation and affinity purification and then identifying Prp19-associated proteins by mass spectrometry. This approach yielded nine core subunits of the complex: Prp19, Cef1, Snt309, Prp46, Syf1, Syf2, Syf3/Clf1, Isy1 and Ntc20 (Ohi et al., 2002; Fabrizio et al., 2009). The human cells have at least three Prp19-like complexes, possibly reflecting the broader involvement of Prp19 in multiple cellular pathways (Chanarat and Sträßer, 2013). In contrast to human, only one NTC/Prp19C has been identified to date in *S. cerevisiae*. Since yeast complex, in addition to splicing, has also been implicated in DNA repair and transcription elongation, we expected existence of multiple Prp19-containing complexes in yeast as well (Chanarat and Sträßer, 2013; Idrissou and Marechal, 2022).

Our findings confirm the presence of a novel Prp19-containing complex in yeast, which appears to have a potentially distinct role in transcription. Specifically, we observed a statistically significant interaction of Prp19 with RNAPII, the general transcription factor TFIID, the CPF 3’ end processing/termination complex, and the RSC chromatin remodeling complex (Fig. 2). Importantly, these interactions were detected exclusively in the context of transcriptionally active chromatin. Furthermore, the interaction of Prp19 with RNAPII and TFIID occurred through direct protein-protein contacts, independent of chromatin DNA or nascent RNA (Table 1A, 1B, 1D and 1E). Although previous studies have implicated yeast NTC/Prp19C in transcription elongation (Chanarat et al., 2011; Henke-Schulz et al., 2024), our results suggest a novel role for Prp19 in the initiation step of transcription. The chromatin-derived Prp19 also exhibited significant interaction with two subunits of Tho complex, which is a part of TREX complex. However, we could not detect any interaction of Prp19 with the previously identified eight core subunits of the spliceosome-derived NTC/Prp19C complex. A possible explanation of these seemingly contradictory results is that Prp19 may associate with different proteins depending on its functional state (Fig. 7). The spliceosome-derived Prp19 interacts with splicing factors, but in the chromatin context, it associates with the factors involved in transcription. Furthermore, all interactions of Prp19 with transcription-linked proteins reported here can withstand high ionic strength (0.5 M ammonium sulfate). It is possible that the interactions of Prp19 with splicing factors are weaker and are disrupted at high salt concentration.

**Figure 7:**
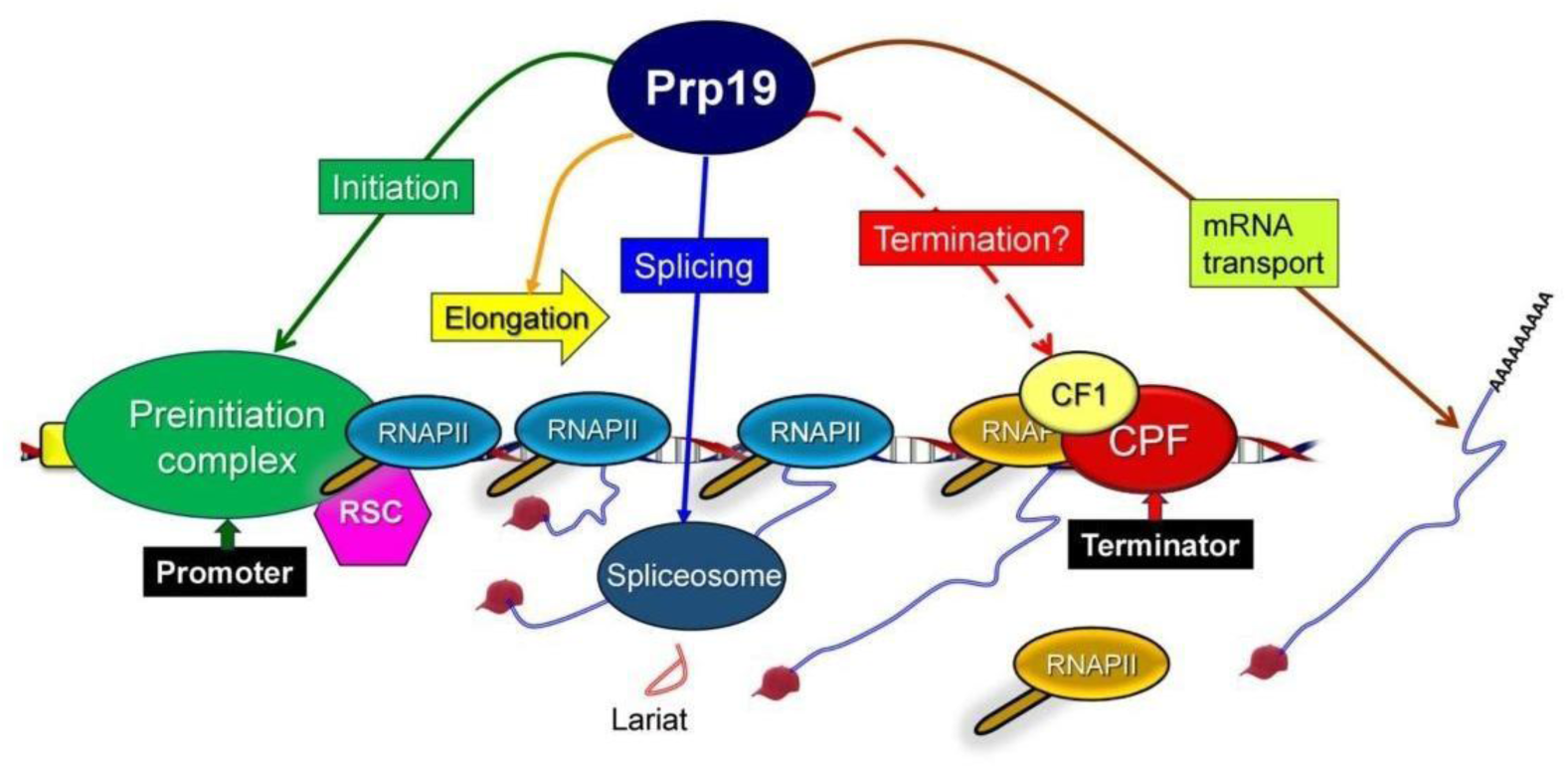
A model showing pleiotropic role of Prp19 in transcription and splicing. Prp19 role in splicing, elongation of transcription and RNA trafficking is already known. Here we show a novel role of Prp19 in the initiation step of transcription. The terminator crosslinking suggest that the protein may have a role in termination as well.

NTC/Prp19C is known to be involved in various aspects of mRNA metabolism, including splicing, transcription elongation, and mRNA export in both yeast and higher eukaryotes (Chanarat and Sträßer, 2013; Idrissou and Marechal, 2022). While its role in transcription elongation has been well established, here, we present evidence for its novel involvement in transcription initiation. Although a direct role of Prp19 in transcription has not been previously demonstrated, the NTC/Prp19C has been implicated in transcription in yeast. Syf1 and Syf2, two subunits of NTC/Prp19C, have been linked to the elongation step of transcription (Chanarat et al., 2011; Henke-Shulz et al., 2024). Here we provide multiple evidence for an additional, previously unrecognized role of Prp19 in initiation of transcription of at least a subset of both intron-containing and intron-lacking genes. First, Prp19 directly interacts with the TFIID and the RSC complex exclusively in the context of transcriptionally active chromatin (Fig. 2). Second, auxin-mediated depletion of Prp19 from yeast cells adversely affected the transcription of a subset of both the intron-containing as well as intron-lacking genes (Figs. 3 and 4). Third, the recruitment of TFIID to promoters of selected genes was severely compromised in the absence of Prp19 (Fig.5). Fourth, Prp19 crosslinked to the promoter, coding region, and the 3’ end of selected genes, with a slight peak at the promoter region (Fig. 6). Given that Prp19 affected transcription of a subset of both intron-containing and intron-lacking genes, we propose that it plays a broader role in transcription and RNA processing. Prp19, however, is not an essential transcription factor as it is required for transcription of only a subset of genes, and transcription of the affected genes was reduced but was not completely abrogated in the Prp19-depleted cells. The interaction with the components of TREX complex corroborates its involvement in transcription-coupled RNA trafficking out of the nucleus (Chanarat et al., 2011). Our analysis also revealed a subset of genes that exhibited enhanced transcription in the absence of Prp19, suggesting that Prp19 has an inhibitory influence as well on transcription. Whether Prp19 is directly or indirectly inhibiting transcription of these genes needs further investigation.

Interestingly, in this new context for transcription initiation, we did not observe Prp19 interactions with the NTC/Prp19 complex. This suggests that the role of Prp19 in transcription initiation is distinct from its role in splicing. Given its extensive involvement in various stages of gene expression, we propose that Prp19 may serve as a crucial integrative hub or scaffolding protein, linking transcription initiation and elongation with post-transcriptional processes such as mRNA export and degradation (Fig. 7).

Our findings open new avenues for research into the multifunctionality of splicing factors. It challenges the traditional view of transcriptional regulation and raises questions about how splicing and transcription are coordinated at the molecular level. Further investigations into the specific molecular interactions between Prp19 and transcription factors will be essential to fully understand the breadth of its regulatory functions. This work underscores the evolving complexity of gene expression regulation and highlights the importance of studying these “moonlighting” factors, which play roles in multiple, seemingly unrelated biological processes.

In conclusion, our study highlights a novel transcription initiation-specific function for the Prp19 complex in yeast, further expanding its already broad functional repertoire. This work contributes to a growing body of evidence that challenges the traditional view of splicing and transcription as separate processes, instead suggesting that factors like Prp19 can bridge these stages to coordinate efficient gene expression (Fig. 7). Further research into the dynamic roles of Prp19 and related complexes will likely reveal new insights into the integration of transcriptional and post-transcriptional regulation.

## Materials and Methods

### Cell culture

A 5 ml culture was started in yeast-peptone-dextrose (YPD) medium using colonies from a freshly streaked plate. The culture was grown overnight at 30°C with constant shaking at 250 rpm. All *Saccharomyces cerevisiae* cell cultures were grown in YPD medium unless otherwise stated. All strains were grown at 30°C and 250 rpm. The next morning, cultures grown overnight were diluted 1:100 in YPD broth and allowed to grow at 30°C with constant shaking until the desired *A_600_* was reached. The cells were then processed accordingly for each experiment.

### Cell culture for auxin-induced degradation of Prp19

The starter culture of Prp19-AID-tagged strain was grown in 5 mL of YPD medium and grown overnight at 30°C and 250 rpm in an orbital shaker. Next morning, the overnight grown culture was diluted 1:100 in appropriate medium and allowed to grow till A600 reached 0.3–0.8. At this point, one-half of the culture was treated with auxin (final concentration 500 μM) and the other with DMSO (control). Both cultures were grown for another 90 minutes. Cells were collected by centrifugation and used for RT-PCR, TRO or ChIP analysis.

### Auxin plating assay

Cells were grown overnight in a 5 mL liquid YPD medium as previously described. Next day, the 5 mL culture was diluted 1:100 in YPD, and cells were allowed to grow until *A_600_* reached ∼0.4-0.6. An aliquot of cell culture was transferred to a well of 96-well plate and serially diluted (1:10, 1:100, 1:1,000, 1:10,000, 1:100,000). The dilution procedure was performed for wildtype cells and cells harboring AID-tagged Prp19. The serially diluted cells were plated on YPD or YPD+500 μM auxin plates using a sterile prong frogger. The plates were incubated at 30°C. Images of plates were taken each day until the growing colonies reached saturation.

### Purification of termination complexes

The CPF, CF1 and Rat1 complexes were purified following the protocol shown in Fig. 1A from strains harboring Myc-tagged Ssu72, HA-tagged Rna15 and HA-tagged Rat1 respectively (Fig. 1B). Yeast cells from 10 L of exponentially growing liquid culture were lysed, and the lysate was separated into soluble and chromatin fractions by differential centrifugation as described in O’Brien and Ansari (2024). The chromatin and soluble fractions were subjected to affinity chromatography on anti-HA or anti-MYC magnetic beads. 100 μL of the bead slurry was transferred to a 1.5 ml microcentrifuge tube and placed on a magnetic rack for 30 seconds to allow for the beads to settle along the magnet. The supernatant was removed, and the beads were washed with wash buffer [25 mM tris-acetate pH 7.8, 5 mM DTT, 1 mM MgCl2, 100 mM potassium acetate, and 0.05% Triton X-100] three times to allow equilibration of beads. The chromatin or soluble fraction was added to the buffer equilibrated beads and mixed gently. Protein solution (chromatin or soluble fraction) was allowed to bind to the affinity beads for 3 hours at 4°C. Binding was performed by gentle shaking the bead and protein solution on a nutator. After binding, the supernatant was carefully removed, and beads were washed three times with wash buffer. Bound proteins were eluted with 250 μL of elution buffer (Tris-HCl pH 6.8, 60 mM, 10% glycerol, 2% SDS and 500 mM β-mercaptoethanol). Elution was performed at room temperature for 30 minutes on a benchtop nutator. The eluted proteins were separated from the beads by placing the tubes on a magnetic rack. The eluent was transferred to a new tube and stored at −80°C.

### MNase Digestion

Micrococcal Nuclease (MNase) digests both DNA and RNA. MNase digestion was performed before affinity purification. The chromatin eluted samples were digested with 20,000 units of MNase in the presence of 5 mM calcium chloride in 1 mL of sample volume at 37°C for 30 minutes.

### Proteomic Analysis

Analysis of the mass spectrometry data from purified protein preparations was done essentially as described in O’Brien and Ansari (2024). The mass spectrometry data were compiled into Scaffold files. The Scaffold program display was set to protein name and species (*S. cerevisiae*), UniProt accession number, alternate protein name identification, molecular weight, and, importantly, the normalized total spectra (spectral counts). Spectral counts greater than 95% were included in our analysis. The protein threshold was set to 0.1% false discovery rate (FDR), the minimum number of peptides was set to 1, and the peptide threshold was set to 0.1% FDR. Spectral counts for each protein in tagged and untagged/control replicate samples were divided by their molecular weight to produce the spectral abundance factor (SAF), as described in Paoletti et al., (2006) and Zybailov et al., (2006). The average SAF value for untagged replicates was subtracted from each tagged replicate SAF value. The SAF values were then normalized against the SAF of the bait/tagged proteins Ssu72, Rna15, and Rat1 to generate the normalized spectral abundance factor (NSAF). Finally, the NSAF values from three biological replicates were averaged to generate a mean NSAF value for each interactor. Following this protocol, the NSAF values of each Ssu72, Rna15 and Rat1 interacting protein in the soluble and chromatin fraction were calculated and tested for significant enrichment by a two-tailed standard t-test. A p-value of 0.05 or less indicated a significant difference between fractions or samples, with standard error accounting for the variability across replicates. The authenticity of the soluble and chromatin fractions was verified using marker proteins, α-tubulin for soluble fraction, and histones for the chromatin fraction of the cell lysate. The mass spectrometry proteomics data have been deposited to the ProteomeXchange Consortium via the PRIDE [1] partner repository with the dataset identifier PXD061187 and 10.6019/PXD061187.

### Transcription run-on (TRO) assay

The overnight grown cultures were diluted to 1:100 in 100 mL YPD and grown to an *A_600_* of 0.8 as described above. The 4-thiouracil was now added to the culture from 2 M stock to a final concentration of 5 mM. The incorporation of 4-thiouracil into nascent RNA was allowed for exactly 5 minutes at 30°C in an orbital shaker. The cell culture was then transferred to 50 mL conical tubes and centrifuged at 3000xg for 2 mins at 4°C. The cell pellet was washed with 20 mL of cold 1xPBS, and washed cell pellet is transferred to a 1.5 mL microfuge tube. The cell pellet was flash frozen with liquid nitrogen and stored in a −80°C freezer. At this stage, samples can be stored overnight, and the protocol is resumed next day.

Next day, samples were removed from storage and thawed on ice. Cells were spun down in a microcentrifuge at 1,100xg for 5 minutes at 4°C. The cell pellet is quickly resuspended in 500 μL of phenol (pH 4.5). An equal volume of AES (50 mM NaOAc pH 5.3, 10 mM EDTA pH 8, and 1% SDS) buffer is added, and the sample is incubated in a 65°C water bath for 5 minutes. During the 5-minute incubation period the sample contents are vortexed for 10 seconds once every minute. The samples were then incubated on ice for 5 minutes followed by addition of 200 μL of chloroform to each sample. The samples were mixed vigorously on a vortexer for 30 seconds, incubated at room temperature for 2 minutes followed by centrifugation at 14,220xg for 5 minutes at 4 °C in a microcentrifuge. The upper aqueous phase was transferred to a new tube and ethanol precipitated for 10 minutes in the presence of 2 μL GlycoBlue before spinning down precipitated RNA. The RNA pellet was washed with 750 μL of ice-cold 75% ethanol and centrifuged at 14,220 x g for 5 minutes at 4 °C. The supernatant was carefully removed, and the pellet was air dried for 5–10 minutes at room temperature. It was important not to let the RNA pellet dry completely as this will greatly decrease its solubility in the next step. Finally, the RNA pellet was resuspended in 100 μL DEPC-H_2_O.

The concentration of RNA was measured using a Nanodrop. The total concentration of RNA for the following steps was adjusted to 2 mg/ml and aliquots of 200 μg of total RNA were made for the remaining processing steps. The 200 μg RNA aliquots were heated for 10 minutes in the 65°C water bath to remove any secondary RNA structures. RNA was then biotinylated by adding 650 μL of DEPC water, 100 μL of the biotinylation buffer (100 mM Tris-HCl pH 7.5 and 10 mM EDTA) and 150 μL of HPDP-biotin (stock 1 mg/ml). The RNA solution was mixed thoroughly and incubated at room temperature in dark for three hours on the nutator in a 2 mL tube. After incubation, an equal volume of chloroform was added to the tube and mixed thoroughly by a vortexer before spinning at 13,000 x g for 5 mins in a 4°C microcentrifuge. The supernatant was transferred to a new 1.5 mL microfuge tube and ethanol precipitated using Glycol-blue as described above. The pellet was resuspended in 100 μL of DEPC-treated water. The RNA suspension is then purified using a Qiagen RNeasy kit following manufacturer’s protocol. The final volume following purification over the RNeasy kit was 200 μL. The concentration of RNA was measured using Nanodrop. The yield of RNA at this stage is in the range of 100-500 μg. RNA at this stage can be stored at −80 °C for several months.

The RNA sample was removed from storage and thawed on ice. The following buffers were added in this order: 25μL 10X NaTM buffer (0.1M Tris-HCl pH 7.0, 2 M NaCl, and 250 mM MgCl2), 25μL NaPi buffer pH 6.8 (0.5 M NaH2PO4 and 0.5M NaHPO4), and 2.5 μL of 10% SDS. The contents were mixed thoroughly and spun down quickly to remove liquid from the sides. In a new 1.5 mL tube, 100 μL slurry of streptavidin beads was added and placed on a magnetic rack. The storage buffer was carefully removed from the magnetic beads. The streptavidin beads were washed with 200μL of bead buffer (200 μL NaTM buffer, 200 μL NaPi buffer, 20 μL 10%SDS, and 1.58 mL ddH2O) followed by a brief vortexing and spinning at max speed for 5 seconds. The tube was placed on the magnetic rack to allow beads to settle to the magnet before removing the buffer. The beads were then blocked by adding 200 μL of bead buffer, 10 μL 20mg/mL glycogen, and 1.25 μL 10 mg/mL E. Coli tRNA and placed on the nutator at room temperature for 20 minutes. The tubes were then placed on the magnetic rack to remove the blocking buffer and washed once with bead buffer as previously described. The RNA suspension was added to the beads after heating at 65°C for 10 minutes in a water bath. After combining the RNA suspension with the magnetic streptavidin beads, the sample was incubated at room temperature on a nutator for 90 minutes to enable bonding of biotinylated RNA to streptavidin beads. The tube was placed on a magnetic rack and the supernatant was removed. The beads were then washed three times with the bead buffer as previously described to remove any unbound RNA. To elute RNA from the streptavidin beads, 0.5 mL of Trizol was added to the tube. The contents were mixed thoroughly and incubated at room temperature for 5 minutes. 100 μL of chloroform was added to the tube and mixed vigorously on a vortexer before being incubated at room temperature for 10 minutes. A chloroform extraction is performed by centrifugation in a tabletop centrifuge at 14,220 x g for 10 minutes at 4 °C. The upper aqueous phase containing RNA is carefully transferred to a 1.5 mL microcentrifuge tube. The RNA is then ethanol precipitated in the presence of 2μL of glycoblue. The supernatant is removed, and the RNA pellet should be visible at the bottom of the tube. It was crucial at this stage to carefully remove any residual buffer as the RNA pellet is flimsy and easily detaches from the wall of the microcentrifuge tube. The pellet was washed with 1 mL of ice-cold 75% ethanol. The supernatant was removed following the wash step and the RNA pellet was air-dried for 5-10 minutes at room temperature. Finally, the pellet was resuspended in 22 μL of DEPC-H_2_O and was ready to be used for subsequent RT-PCR or GRO-seq analysis.

### GRO-Seq

GRO-seq was performed essentially as described in O’Brien et al., (2024) except that 4-thiouracil was used instead of Br-dUTP. 4-thiouracil labeling was performed as described in TRO assay above. Nascent, isolated RNA obtained using GRO-Seq was obtained from three biological replicates.

### GRO-Seq analysis

Raw GRO-Seq reads were first assessed for quality using FastQC and were then aligned to the *Saccharomyces cerevisiae* S288C genome (SGD version R64-1-1) using STAR (v2.7.11a) with the following parameters: maximum intron size of 2,000 nucleotides, a minimum overlap for paired-end alignment of 10 and sorted BAM alignment format. Files were then sorted by co-ordinates using STAR. Alignment quality was evaluated using SAM tools (v1.9), and BAM files were indexed for downstream analysis.

Count matrix was generated using featuresCount–countReadPairs -s2 -T6 -texon (subread-2.06). The counts obtained were processed in R/Bioconductor as described in Gentleman et al., (2004) and Ihaka and Gentleman (1996). Differential expression analysis was performed using DESeq2. It identified significantly differentially expressed genes with an adjusted p-value < 0.05 and a log₂ fold-change of greater than 2.

Bamcoverage (version) was used to generate bigwig files, which were imported into Integrated Genome Browser (IGV) for gene browser track visualization. The subsequent data in the bigwig file, representing nascent transcription levels from GRO-seq, was selectively enriched for the genomic regions specified in the BED files using the computeMatrix function (Galaxy Version 3.3.0.0.0). A heatmap was generated using plotHeatmap (Galaxy Version 3.3.0.0.1) to visualize changes in transcriptional activity across these regions under wild-type conditions and Prp19-depleted conditions (+auxin). Additionally, plotProfile (Galaxy Version 3.3.0.0.0) was used to generate metagene plots, allowing for the comparative analysis of nascent RNA distributions across annotated genomic regions.

The following BED files were used in this study:

1. 93 intron-containing genes with transcription start site (TSS) and transcription end site (TES) coordinates.
2. 302 non-intronic genes with TSS and TES coordinates.

Genes were classified as TATA-containing or TATA-less based on a curated dataset of yeast genes characterized by Basehoar et al., (2004). Gene ontology analysis was performed using clusterProfiler with org.Sc.sgd.db annotation.

### Chromatin Immunoprecipitation (ChIP)

Cells were grown in appropriate medium as described above in 100 mL culture. Once the cells reached an *A_600_* ∼ 0.7-0.8 crosslinking was performed with 1% formaldehyde (2.7 ml stock/100 mL cell culture) for 20 minutes at room temperature with vigorous shaking. The reaction was stopped by addition of glycine to a final concentration 125 mM, and cultures were shaken vigorously for an additional 5 minutes at room temperature. The cell culture was transferred to a 50 mL tube and spun for 5 minutes at 3,000 rpm in a Sorvall RC6Plus centrifuge. The supernatant was discarded, and the cell pellet was washed once with 10 mL of ice-cold 1xTBS buffer containing 1% Triton X-100, and twice with 1xTBS buffer only. The cell pellet was resuspended in 800 μL of cold FA-lysis buffer (50 mM HEPES-KOH pH 7.9, 140 mM NaCl, 1 mM EDTA, 1% Triton X, 0.1% sodium deoxycholate, 1 mM PMSF, and 0.1% SDS). The cell suspension was then frozen with liquid nitrogen and stored at −80°C.

Next, cells were thawed and approximately 400 μL of acid-washed glass beads were added to each tube. Cells were lysed by vigorous shaking at 4°C for 40 minutes. The cell lysate was collected by puncturing the bottom of the tube with a red hot 22-gauge needle and collecting the lysate in a 15 mL tube by spinning at 1,000 rpm at 4°C in a Sorvall RC6Plus centrifuge. The filtrate was then transferred into a 1.5 ml microfuge tube and spun at 4°C for 15 minutes at the maximum speed in a microcentrifuge. The supernatant was discarded, and the crude chromatin pellet was washed with 1,000 μL of FA-lysis buffer and resuspended in 1,000 μL of FA-lysis buffer. The crude chromatin preparation obtained was diluted to 4 mL by the addition of FA-lysis buffer and 40 μL of PMSF. Chromatin was sheared by sonication with thirty to forty-five pulses (depending on gene size) of 20 seconds each with 30 second cooling after each pulse (This will result in a total sonication time of 10-15 minutes). Sonication was performed at the 30% duty cycle in a Branson digital sonifier. Following sonication, samples were centrifuged at 14,000 rpm for 15 minutes in a 4°C centrifuge. The pellet was discarded, and the supernatant was used in subsequent steps. The supernatant can be frozen in liquid nitrogen and stored at −80°C at this stage.

The sonicated chromatin was thawed and 300 μL of chromatin is used for each ChIP assay. 50 μL is stored aside as an input control. Approximately 5-10 μg of appropriate antibody-conjugated magnetic beads were added to a 1.5 mL tube (Pierce anti-c-Myc magnetic beads (88842) for Prp19; Santa Cruz Biotechnology TBP antibody (sc-74596) for TBP ChIP). The storage buffer was removed by a brief spin and 300 μL of chromatin was added to the beads. The chromatin was allowed to bind to beads for 4 hours at 4°C with gentle shaking on a nutator. The beads were washed successively with 1 mL each of FA-lysis buffer containing 0.25% SDS, FA-lysis buffer containing 500 mM NaCl and 0.25% SDS (50 mM HEPES-KOH pH 7.9, 500 mM NaCl, 1 mM EDTA, 1% Triton X, 0.1% sodium deoxycholate, 1 mM PMSF, and 0.25% SDS), ChIP wash buffer 0.25% SDS (10 mM Tris-HCl pH 7.5, 250 mM LiCl, 0.5% Triton X, 1 mM EDTA pH 8, 0.5% sodium deoxycholate, and 0.25% SDS) and 1XTE buffer. The washing steps were performed twice at room temperature with the 1.5 mL tubes being placed over the magnetic rack to remove the supernatant containing unbound chromatin. After washing, the beads were resuspended in 100 μL of ChIP elution buffer (50 mM Tris-HCl pH 7.5, 1% SDS, and 10 mM EDTA pH 8) and incubated at 65°C for 10 minutes in a water bath to elute chromatin from the magnetic beads. The 1.5 mL tubes containing the chromatin-bead mix was placed over the magnetic rack to remove the supernatant to transfer it to a new 1.5 mL tube. The elution was performed twice. There should be 200 μL of eluted chromatin after the elution steps. The chromatin eluent was incubated with 10 μg of DNase-free RNase (Worthington) for 30 minutes in a 37°C incubator. 20 μg of proteinase K and 2.5 μL10% SDS were added and incubated at 42°C for 90 minutes in a water bath. Finally, the protein-DNA crosslinks were reversed by overnight incubation at 65°C in a water bath.

Next day, samples were extracted with equal volumes of DNA phenol-chloroform at least two times followed by ethanol precipitation of DNA in the presence of carrier glycogen. The DNA pellet was resuspended in 50 μL of 1XTE and used as template for further PCR analysis.

## Data Availability

All relevant data are within the paper and its [Supporting information](https://journals.plos.org/plosbiology/article?id=10.1371/journal.pbio.3002848#sec032) files. The mass spectrometry proteomics data have been deposited to the ProteomeXchange Consortium via the PRIDE partner repository with the dataset identifier PXD061187 and 10.6019/PXD061187. Statistical source data is provided with this article. The GRO-Seq data has been deposited in NCBI database. The NCBI Geo accession number is: GSEXXXXX. All other relevant data that support this study is available from the corresponding author upon reasonable request. All yeast strains are available upon request.

## Acknowledgements

We thank Dr. Markus Friedrich of WSU for critically reading the manuscript and for sharing valuable comments. We thank Dr. Marc Gartenberg for kindly providing plasmids for making Prp19-AID strain. We thank lab members Emma Fidler, Kasun Rathnasinghe, Isabella Nadeau and Kyle Kilgore for useful help. We thank DNA Core facility of WSU for sequencing GRO-Seq library. We acknowledge the assistance of Dr. Paul Stemmer of Wayne State University Proteomics Core that is supported through National Institutes of Health grants P30 ES020957, P30 CA 022453 and S10 OD010700.

## Author Contributions

AA (Athar Ansari) conceptualized, supervised research and wrote the manuscript. KD performed all experiments and data analyses. MAE helped in protein purification and proteomic data analysis. AA (Ahlam Awada) performed TBP ChIP. ZD helped in GRO-Seq analysis. All authors approve the submitted version.

