## Supporting Information for "Protein-interaction network analysis reveals a role of Prp19 splicing factor in transcription of both intron-containing and intron-lacking genes"

### Supporting Table 1: Yeast strains

| Strain | Genotype | Reference |
|---|---|---|
| BY4733 | MATa his3Δ200 trp1Δ63 leu2Δ0 met15Δ0 ura3Δ0 | This study |
| H98 | MATa ura3-52 trp1Δ63 leu2Δ1 | This study |
| WA89 | MATa ura3-52 trp1Δ63 leu2Δ1; Rna15-HA tag | This study |
| WA322 | MATa ura3-52 trp1Δ63 leu2Δ1; Rat1-MYC tag | This study |
| WA326 | MATa ura3-52 trp1Δ63 leu2Δ1; Ssu72-MYC tag | This study |
| WA491 | MATa his3Δ200 trp1Δ63 leu2Δ0 met15Δ0 ura3Δ0; Prp19-MYC tag | This study |
| WA470 | MATa his3Δ200 trp1Δ63 leu2Δ0 met15Δ0 ura3Δ0; Prp43-MYC tag | This study |
| WA493 | MATalpha ura3::pADH1-OsTIR1-9MYC::URA3 Prp19-AID tag | This study |

### Supporting Table 2: Primes

#### Yeast strain construction primers

| Name | Sequence |
|---|---|
| 5' PRP19-MYC | CCGCTATTCTGAAGACAAATGATAGTTTCAATATTGTTGCATTGACACCCCGGATCCCCGGGTTAATTAA |
| 3' PRP19-MYC | TTACTACTATTACACAGGTTTATTTAGAAAGTACAAACGTGTCAGCGTATGAATTCGAGCTCGTTTAAAC |
| 5' PRP43-MYC | ACGAGTTGAAACAAGGTAAAAACAAAAAGAAGAGTAAGCACTCCAAGAAACGGATCCCCGGGTTAATTAA |
| 3' PRP43-MYC | CCTATAAATTTATATAAATCTATTTTTTTTTTTTTTTCGACACAAAATGTTGAATTCGAGCTCGTTTAAAC |
| 5' PRP19-AID | CCGCTATTCTGAAGACAAATGATAGTTTCAATATTGTTGCATTGACACCCCGTACGCTGCAGGTCGAC |
| 3' PRP19-AID | TTACTACTATTACACAGGTTTATTTAGAAAGTACAAACGTGTCAGCGTATTCGATGAATTCGAGCTCG |
| 5' PRP19-MYC | CCGCTATTCTGAAGACAAATGATAGTTTCAATATTGTTGCATTGACACCCCGGATCCCCGGGTTAATTAA |

**ChIP primers**

***APE2***

| Name | Sequence |
|---|---|
| 5' CHIP | GGTCCTGTTGTATATGACTTTGT |
| 3' CHIP | AAAGAATACTGTTGATCTGCCC |
| 5' CH2P | CAAAATAATCAACTGTAGTGGTTCA |
| 3' CH2P | GCTCGTTGACAAGACCCTT |
| 5' A | CCAATTGTTCGGTGGCTATT |
| 3' A | ACAAATCTTGGGGGAGTTAATT |
| 5' B | TGAAGTACGTTGAATCCAAG |
| 3' B | TTCGTGAACAGCAACATTG |
| 5' EF1 | AGTTTGGGAACAGTATGTTACTG |
| 3' EF1 | TCACAGAACGAACGTCTTTAC |
| 5' C | ACTATCTGCTTCTGGCTAC |
| 3' C2 | AAACATGGTCACTTTCAAACGT |
| 5' T4 | GTGGTTGCAAACATACATAAATC |
| 3' T4 | TACGATGAAGTTGCTTGTTATCG |
| 5' CHIT | CAACTGGACCAGTGAATAGAAC |
| 3' CHIT | CCTTTTTTCTCCTCATCTCGGT |

***ASC1***

| Name | Sequence |
|---|---|
| 5' UP DIAG | ACTGGTAATACTCATCACCATACTTATAT |
| 3' DN3 | CACAATTAAAGGAATAGCCCAA |
| 5' UP3 | TTTTTGGCATTGGGCTATTC |
| 3' DN2 | CACTTTATTTACTTTAGTGTGTTATAAGG |
| 5' P2 | GACTGCTCCTTTGGTTTTCC |
| 3' P2 | GGTTGACCAGCAGAAGTAGCC |
| 5' A | CTTACGCTTTGTCTGCTTCTTG |
| 3' A | GATGGTCTTGTCACGGGAAC |
| 5' R | TTGGCTGCTAAGAAGGCTATG |
| 3' C | CCAAGCCAAAGAAACAGCAT |
| 5' CCC T1 | CCGGTTACACCGACAACG |
| 3' T3 | GCCAAGGAGACTGAATTTAATG |
| 5' T1 | TAATATAATCGTCATAGATTTCGAAG |
| 3' T2 | CTATGGAATGGGGGTTTTAAG |
| 5' S | TTATGTATCTTCTAGTTATTGGTCATG |

***IMD4***

| Name | Sequence |
|---|---|
| 5' P | ATGATAGACTTGAGGGACGCAC |
| 3' P4 | CAGCATCTCGAAGAGCAAAAAA |
| 5' P2 | GCCATATAAATATCAGTTGAGAATCC |
| 3' P2 | GTATGTCTTCAAATGTTCTAAAGCC |

| | |
|---|---|
| 5' CCC | TGGATTACAAAAAGGCTTTAGAAC |
| 3' CCC | CTTCAGTGACTGTGTCCATAGGAG |
| 5' B | AGTTTGGTTTCTCTGGCTTCC |
| 3' B | GATCACAGTCAGTAGAACGAAAGTT |
| 5' D | AACTTTCGTTCTACTGACTGTGATC |
| 3' D | CTCAACTTTATGAAGACTGAAGAAAT |
| 5' T2 | GAGTAAGCATCCATAGATATTTAAAAG |
| 3' T2 | CGAACTGAAAAACGAAAATAAGAA |
| 5' T1 | CAAAACTAAAATAAAGAACAGACC |
| 3' T | GGTGTAAAGGGTTTAGATGAAGC |

***BUD3***

| **Name** | **Sequence** |
|---|---|
| 5' P0 | AATCTCTGGTGGATTTTTACCG |
| 3' P2 | GCTTAGGGTTCTTTTTTGTC |
| 5' P | GTGGTGCTCTTTGTCATACGC |
| 3' P | TAATGAGTAGCAGGTAGCAGCAG |
| 5' H2 | TACTACACTTCAGATGGATCAAACAG |
| 3' H2 | TGACGAAGAAGAGATAAATGCAG |
| 5' I | CGCAGAAACTCTCTCCACAAG |
| 3' I | CGGCGCTAACCTAGGAATAG |
| 5' ter | TGCTGTTAACTAAAGATGCTTCG |
| 3' ter | TTATTTGTCCATATTTTATGTAAATGT |
| 5' T | GCAATGTATACATTTACATAAAATAT |
| 3' T | GATAATGTAGATTTCGATAAGGCTAG |
| 5' T3 | GATGAAGATGGGAAACAGAACTG |
| 3' T3 | TTGATAGTAGGTAAAGGTATCTAATAT |

***HEM3***

| **Name** | **Sequence** |
|---|---|
| 5' P4 | ACCATCTTTACGCAGCAATTC |
| 3' P6 | CTAATACAGCCCTGTACCATG |
| 5' P5 | CTTCTCTCCTTGTCAATTGAC |
| 3' P5 | GTCAATCTGCCAATGACCTTG |
| 5' P6 | CCGATGAGTAAGCAATGAAGG |
| 3' P7 | ATACGGGATGAGAAGAACAG |
| 5' A | GGTGGGAGAAAATCGAAATTG |
| 3' A | CATGACAAGACAATCTGTTGG |
| 5' C | CGATCCAACAGATTGTCTTG |
| 3' C | GAATCTTCATCATCTTGGTGTC |
| 5' M | AGAAAGGGTGACACCAAGATG |
| 3' M | TTCTGTGCCTTCAACGTCAAC |
| 5' T5 | CACCATTAATTTGATCAGTCC |
| 3' T4 | ATATTTATGCAGTGGTCATTTG |
| 5' T6 | CCTTATCTCCTTTATCTTTCAGT |

| | |
|---|---|
| 3' T6 | TTTTTAGAAGTTCACCAAGTG |

***SUR1***

| Name | Sequence |
|---|---|
| 5' A1 | GTTGTGTATTGACGATACTG |
| 3' A1 | ATCGATGTATACACCACC |
| 5' A2 | ATGTCTCGATCTACATCC |
| 3' A2 | ACGCAGGATAAGATGTGG |
| 5' F2 | GCGCATACCTAAGAACG |
| 5' F0.5 | CCACATCTTATCCTGCG |
| 3' R1 | GCAGTATCACTGGTACTT |
| 5' F0 | GCTGGTTGTGTTCCAAG |
| 3' R2 | CCAGTGAGTACTTAGAAG |
| 5' DN1 | CTCCAACAATTCATATCAGAC |
| 3' R5 | GTAGTATGTCGTAAGAAGC |
| 5' DN3 | CTTCTTACGACATACTACTCC |
| 3' R6 | ATGATTATGTTGATGCTGG |

**TRO primers**

***SUR1***

| Name | Sequence |
|---|---|
| 5' A1 | GTTGTGTATTGACGATACTG |
| 3' A1 | ATCGATGTATACACCACC |

***SEN1***

| Name | Sequence |
|---|---|
| 5' B | CTCATTCAATGAAGAAGCTACG |
| 3' B | TTCGAAAGTAGCCTTAAGTTGC |

***BUD3***

| Name | Sequence |
|---|---|
| 5' H | GACGAATTACCCACAGAATCG |
| 3' H | GGGATTGCTGTCTTTAGCGAG |

***HEM3***

| Name | Sequence |
|---|---|
| 5' B | CTGATCGAAGAAAAGTATCCG |
| 3' B | TGGAAGGTCATCCAGAGAC |

***IMD4***

| Name | Sequence |
|---|---|
| 5' B | AGTTTGGTTTCTCTGGCTTCC |

| | |
|---|---|
| 3' B | GATCACAGTCAGTAGAACGAAAGTTA |

***HPC2***

| **Name** | **Sequence** |
|---|---|
| 5' A | TTTGGATGTTGAAAGCACTGC |
| 3' A | CCGTGCTATTTCCACCTTTAT |

***APE2***

| **Name** | **Sequence** |
|---|---|
| 5' A | CCAATTGTTCGGTGGCTATT |
| 3' A | CACAAATCTTGGGGGAGTTAATT |

***ASC1***

| **Name** | **Sequence** |
|---|---|
| 5' A | CTTACGCTTTGTCTGCTTCTTG |
| 3' A | GATGGTCTTGTCACGGGAAC |

**18S**

| **Name** | **Sequence** |
|---|---|
| 5' 18S | GGAATAATAGAATAGGACGTTGG |
| 3' 18S | GTTAAGGTCTCGTTCGTTCGTTATCG |

**Oligo dT primer**

| **Name** | **Sequence** |
|---|---|
| 3' Oligo dT | TTTTTTTTTTTTTTTTTTTTTTTTT |

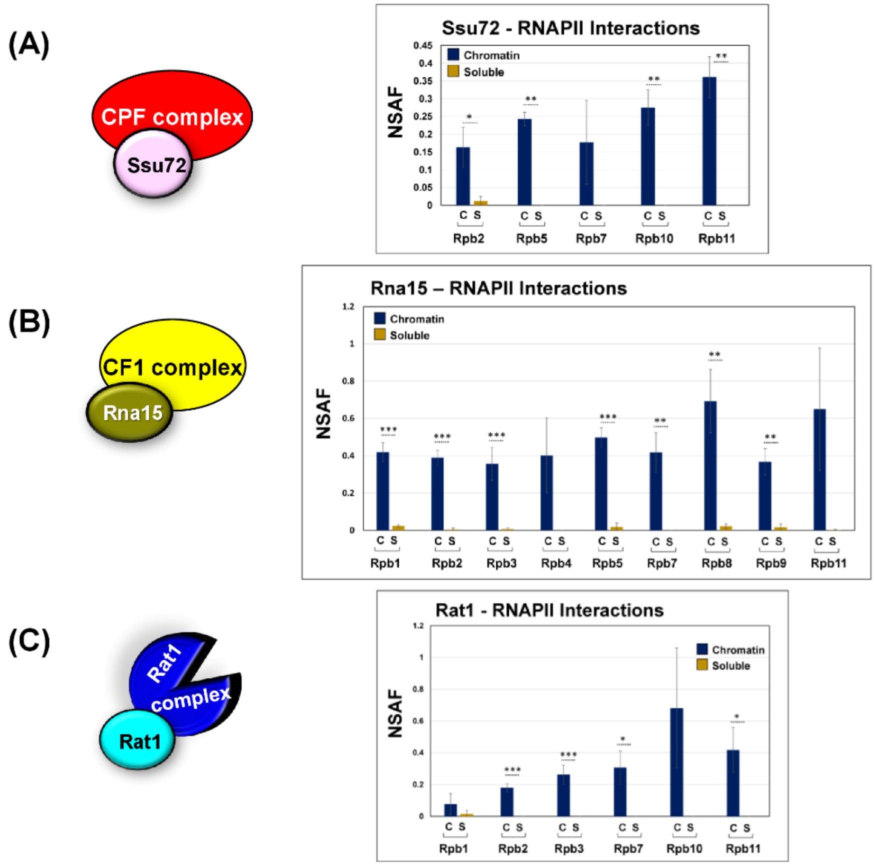


**Supplementary Figure 1:** All three termination complexes in yeast interact with RNAPII subunits in the chromatin fraction. **(A)** Ssu72 subunit of CPF complex, **(B)** Rna15 subunit of CF1 complex, and **(C)** Rat1 subunit of Rat1 complex all associate with multiple subunits of RNAPII in the chromatin context. *p*-values calculated by two tailed t-test indicate significant enrichment of the splicing factors factor in chromatin fraction relative to soluble fraction. One asterisk (*) signifies a *p*-value equal to or smaller than 0.05 ($p \leq 0.05$); two asterisks (**) signify a *p*-value equal to or smaller than 0.01 ($p \leq 0.01$); while three asterisks (***) signify a *p*-value equal to or smaller than 0.01 ($p \leq 0.001$). Error bars represent one unit of standard deviation based on four biological replicates.

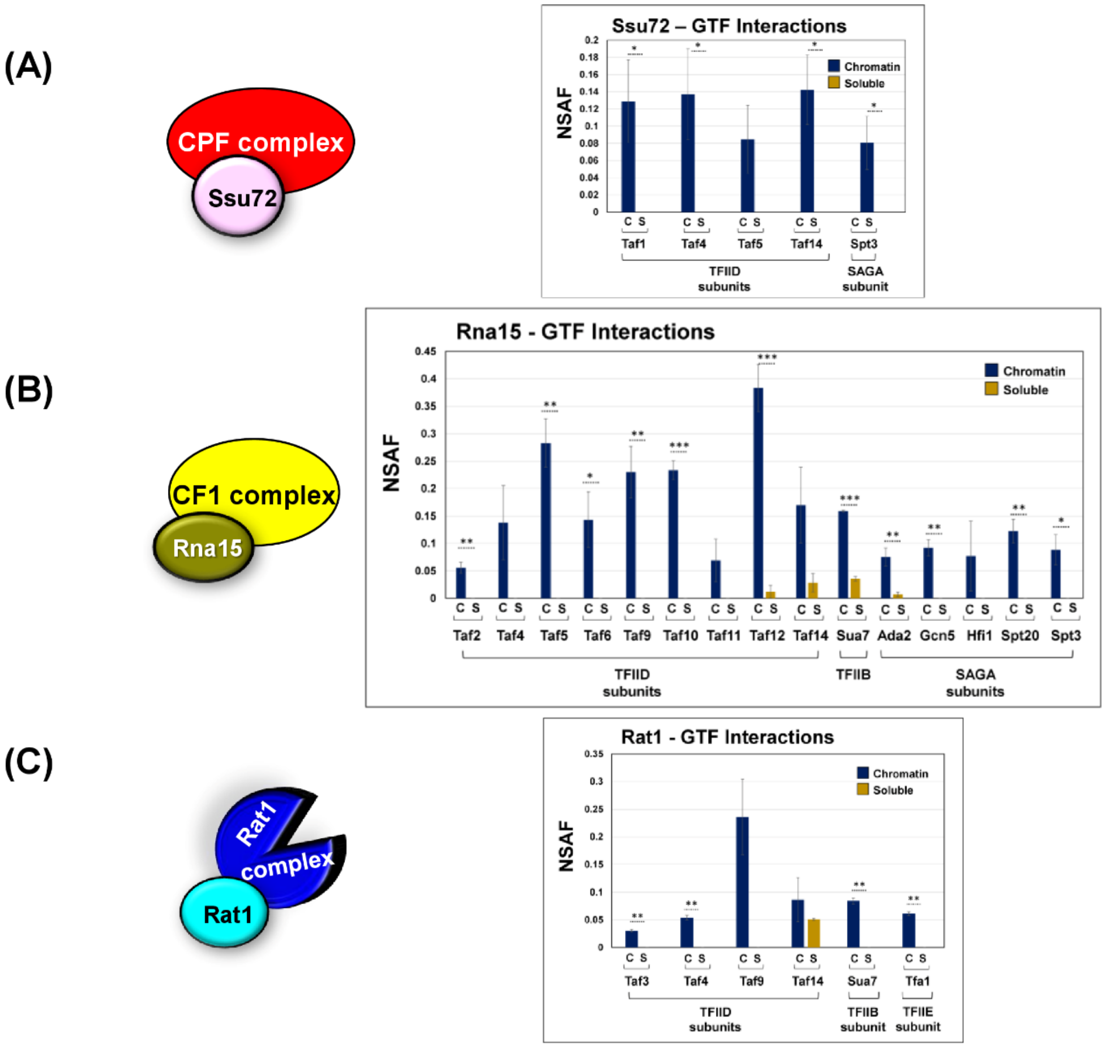


**Supplementary Figure 2:** All three termination complexes in yeast interact with general transcription factors in the chromatin fraction. **(A)** The Ssu72 subunit of CPF complex associates with subunits of TFIID and SAGA complex. **(B)** Rna15 subunit of CF1 complex interacts with TFIID, TFIIB and SAGA subunits. **(C)** Rat1 subunit of Rat1 complex associates with subunits of TFIID, TFIIB, and TFIIE in the chromatin context. *p*-values calculated by two tailed t-test indicate significant enrichment of the splicing factors factor in chromatin fraction relative to soluble fraction. One asterisk (*) signifies a *p*-value equal to or smaller than 0.05 ($p \leq 0.05$); two asterisks (**) signify a *p*-value equal to or smaller than 0.01 ($p \leq 0.01$); while three asterisks (***) signify a *p*-value equal to or smaller than 0.01 ($p \leq 0.001$). Error bars represent one unit of standard deviation based on four biological replicates.

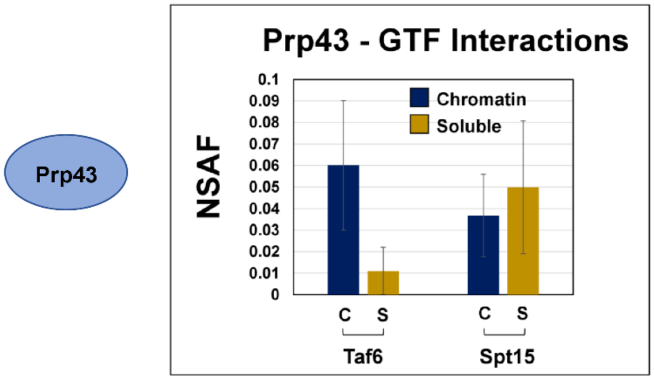


**Supplementary Figure 3:** The splicing factor Prp43 did not display any significant enriched interaction with GTFs in the chromatin fraction. Although Prp43 interacted with Taf6 and Spt15, the protein did not have significant enrichment in the chromatin as compared to soluble fraction. Error bars represent one unit of standard deviation based on four biological replicates.

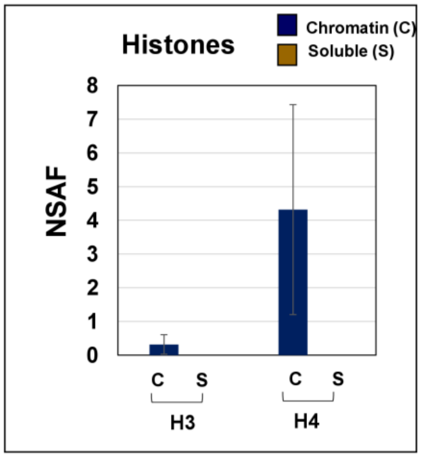


**Supplementary Figure 4:** Validation of the separation of chromatin and soluble fractions. Only the chromatin fraction was enriched with H3 and H4 histone components. Error bars represent one unit of standard deviation based on four biological replicates.

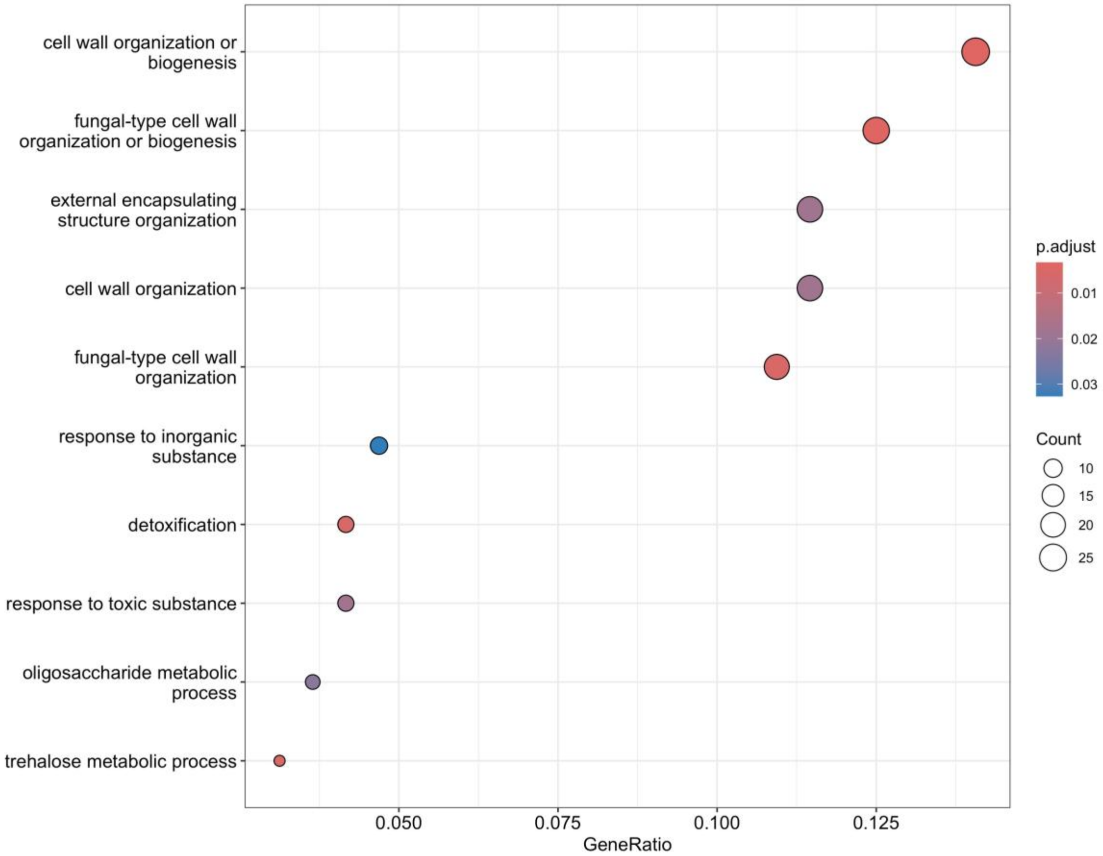


**Supplementary Figure 5:** Ontological analysis revealed that Prp19 has an inhibitory role in transcription of genes linked to cell wall and membrane biology. The plot shows specific categories of genes that are upregulated upon auxin-mediated depletion of Prp19 from yeast cells.
